# Serotonin receptor 4 in mature excitatory hippocampal neurons modulates mood and anxiety

**DOI:** 10.1101/758151

**Authors:** Remzi Karayol, Lucian Medrihan, Jennifer L. Warner-Schmidt, Meghana N. Rao, Eva B. Holzner, Paul Greengard, Nathaniel Heintz, Eric F. Schmidt

**Author notes:** Correspondence (E.F.S.).

## Abstract

Serotonin receptor 4 (5-HT_4_R) plays an important role in regulating mood, anxiety, and cognition, and drugs that activate this receptor have fast-acting antidepressant (AD)-like effects in preclinical models. However, 5-HT_4_R is widely expressed throughout the central nervous system (CNS) and periphery, making it a poor therapeutic target and difficult to pinpoint the cell types and circuits underlying its effects. Therefore, we generated a Cre-dependent 5-HT_4_R knockout mouse line to dissect the function of 5-HT_4_R in specific brain regions and cell types. We show that the loss of functional 5-HT_4_R specifically in the mature excitatory neurons of hippocampus led to robust AD-like behavioral responses and an elevation in baseline anxiety. 5-HT_4_R was necessary to maintain the proper excitability of mature dentate gyrus (DG) granule cells and cell type specific molecular profiling revealed a dysregulation of genes necessary for normal neural function and plasticity in cells lacking 5-HT_4_R. These adaptations were accompanied by an increase in the number of immature neurons in ventral, but not dorsal, dentate gyrus, indicating a broad impact of 5-HT_4_R loss on the local cellular environment. This study is the first to use conditional genetic targeting to demonstrate a direct role for hippocampal 5-HT_4_R signaling in modulating mood and anxiety. Our findings also underscore the need for cell type-based approaches to elucidate the complex action of neuromodulatory systems on distinct neural circuits.

## Introduction

Mood and anxiety are governed by a complex neural circuit comprised of anatomically and functionally distinct components that are distributed throughout the CNS. Functional imaging and postmortem analyses of brains of patients suffering from depression and/or anxiety disorders suggest a dysfunction in regions of the brain that control emotion, including limbic structures such as the hippocampus, amygdala, prefrontal cortex and hypothalamus^1,2^. Each of these areas is strongly modulated by the neurotransmitter serotonin (5-HT) while drugs that target 5-HT signaling, such as selective serotonin reuptake inhibitors (SSRIs), are a first line treatment for affective disorders^3^. The widespread influence of 5-HT on numerous cognitive and behavioral processes leads to many unwanted side effects due to off-target engagement of AD drugs^4^ as well as functional and anatomical separation of the neural substrates underlying pathology from those mediating therapeutic responses^5,6^. This notion is reinforced by regional differences in the expression and regulation of genes and gene networks across the emotion circuit in disease-related contexts^7,8^. Improved treatments for these diseases are dependent on a more precise understanding of cell types and molecular pathways mediating 5-HT-dependent signaling.

The 5-HT_4_ receptor (5-HT_4_R) is one of over 14 known 5-HT receptors in mammals and is strongly linked to AD responses^9^. 5-HT_4_R is an excitatory G_s_-coupled receptor that activates the cyclic AMP (cAMP)-PKA pathway and promotes the excitability of neurons^10,11^. Changes in 5-HT_4_R binding was observed in several brain regions in depressed patients^12,13^ while polymorphisms in *HTR4* (the gene that encodes 5-HT_4_R) were associated with a susceptibility to unipolar depression^14^. In preclinical studies, short-term treatment with 5-HT_4_R agonists, including RS67333, has anxiolytic and antidepressant properties and mimics the cellular and molecular AD responses achieved after chronic SSRI administration^15–19^. In addition, 5-HT_4_R agonists potentiate the effects of SSRIs^20^ while constitutive *Htr4* knockout mice have reduced behavioral responses to acute stress and novelty^21^. 5-HT_4_R has also been linked to the regulation of adult neurogenesis in the dentate gyrus (DG)^15,18^ which is a reliable readout of AD efficacy and may underlie some therapeutic AD effects^22^.

In the brain, 5-HT_4_R is highly expressed in many of the regions linked to mood and anxiety, namely the hippocampus, amygdala, prefrontal cortex, and striatum^10,23^. It is also expressed in peripheral tissues, including the gut, heart, and adrenal glands^24^. Such widespread expression in the CNS and periphery complicates the use of 5-HT_4_R as a viable therapeutic target, as its activation can lead to significant gastrointestinal and cardiac complications^25,26^. To date, most in vivo studies investigating 5-HT_4_R function have relied on systemic pharmacological approaches or constitutive deletion. Therefore, employing a cell type centered approach to elucidate distinct roles for 5-HT_4_R signaling in discrete circuits and/or cell types will facilitate the development of more precise and efficacious therapies by identifying novel targets.

To achieve this, we generated mice harboring a conditional mutant allele to delete functional 5-HT_4_R from genetically targeted cell populations expressing Cre recombinase. We then utilized this line to investigate the role of 5-HT_4_R signaling in excitatory hippocampal neurons on mood since the hippocampus plays a central role in affective behaviors, is highly connected to other brain regions that encode other aspects of emotion^27^, and 5-HT_4_R is expressed at higher levels compared to other brain regions^28^. We report that the loss of 5-HT_4_R specifically from the mature excitatory neurons of the hippocampus led to AD-like effects at the behavioral, cellular, and molecular levels which was accompanied by an anxiogenic phenotype. Cell type specific molecular profiling revealed that the ventral dentate gyrus (vDG) granule cells (GCs) underwent robust molecular adaptations in the absence of 5-HT_4_R. In addition, we observed enhanced neurogenesis in the vDG and reduced excitability of mature vDG GCs in the absence of 5-HT_4_R. Our data reinforce a direct role for hippocampal 5-HT_4_R in mediating affective behaviors, likely through the regulation the excitability of individual neurons and relevant intracellular pathways, and uncovers an unexpected complexity within the hippocampal circuit due to the differential effect on depression- and anxiety-related behaviors.

## Results

### Generation and validation of an Htr4^Floxed^ mouse line

To study the function of the 5-HT_4_R in specific cell types, we generated a novel mouse line (Htr4^Floxed^) to conditionally drive the deletion of functional 5-HT_4_R from genetically defined cell populations using the Cre/loxP system (**Fig. 1a**). Homologous recombination was used to insert loxP sites flanking exon 5 of the mouse *Htr4* gene (**Fig. S1a**). In silico translation analysis predicted that the deletion of exon 5, which encodes the fourth transmembrane domain of 5-HT_4_R, would result in a frame shift mutation and the introduction of stop codons downstream, including one right after the deletion site (**Fig. S1b**), leading to a truncated, non-functional, and/or unstable protein tagged for degradation. This should result in animals harboring a null *Htr4* allele.

**Fig. 1.**
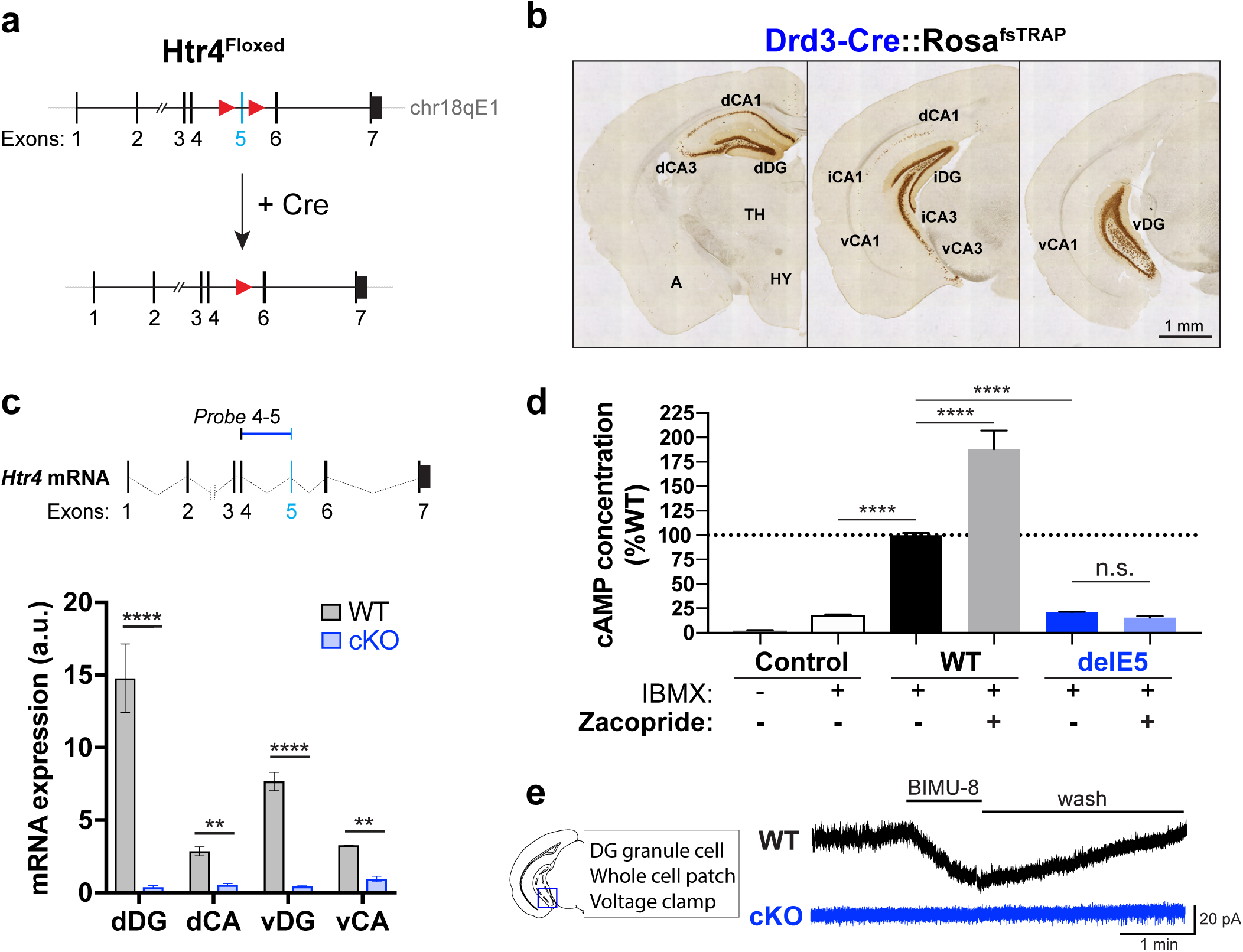
Generation and validation of an Htr4^Floxed^ mouse line. **a**, Schematic of Htr4^Floxed^ allele. LoxP sites (red triangles) were inserted to flank *Htr4* exon 5 (light blue bar) for Cre-mediated excision. **b**, Low magnification anti-EGFP immunohistology of coronal brain sections from a Drd3-Cre::Rosa26^fsTRAP^ mouse showing Cre expression in the hippocampus. dCA, iCA, vCA1/3: dorsal, intermediate, ventral CA1/3 fields, respectively; A: amygdala; TH: thalamus; HY: hypothalamus. **c**, Schematic of the TaqMan probe spanning *Htr4* exons 4 and 5 (Probe 4-5) is shown a top. Below, qRT-PCR (mean ± SEM) quantification showing diminished expression of *Htr4* transcripts containing exon 5 along the dorsoventral axis of the hippocampus in Cre-negative (WT, gray) and cKO (blue) mice. Two-way ANOVA: genotype factor: F(1,24) = 439.0, p < 0.0001 followed by post hoc Fisher’s LSD test. n = 2-6 per group. **d**, Quantification (mean ± SEM) of cAMP induction HEK293T cells expressing EGFP (Control), intact 5-HT_4_R (WT), or *Htr4^delE5^* (del5) in the presence or absence of the 5-HT_4_R agonist, zacopride. One-way ANOVA followed by post hoc Fisher’s LSD test, n = 4 per group. **e**, Representative traces of whole-cell voltage clamp recordings from DG GC in WT and cKO mice with bath administration of 10 mM BIMU-8. Scale bar: 1 min, 20 pA. Fisher’s LSD test, ****p < 0.0001, ***p < 0.001, **p < 0.01, n.s. p > 0.05.

Because *Htr4* is highly expressed in the hippocampal formation^28^, we validated the deletion of exon 5 and the loss of functional 5-HT_4_R by generating a hippocampus-specific *Htr4* conditional knockout. The BAC transgenic Drd3-Cre driver line (founder KI198) was utilized since it expressed Cre recombinase specifically in the hippocampus^29^ (gensat.org; **Fig. 1b**). Homozygous Htr4^Floxed^ mice (Htr4^loxP/loxP^) were crossed to Drd3-Cre mice to generate Drd3-Cre::Htr4^loxP/loxP^ (cKO) mice (**Fig. S1c**). To determine relative *Htr4* transcript levels following Cre-mediated recombination, RNA was extracted from dorsal and ventral subdivisions of the CA fields and DG of cKO mice and Cre-negative littermates (WT; **Fig. S1d**). Quantitative RT-PCR (qRT-PCR) using probes spanning exons 4 and 5 (4-5) of *Htr4* revealed negligible expression of exon 5 in cKO samples compared to WT along the dorsoventral axis of the hippocampus (**Fig. 1c**), while probes spanning exons 3-4 and 6-7 showed comparable expression between genotypes (**Fig. S1e**). These data confirm the generation of a mutant *Htr4* transcript (*Htr4^delE5^*) following Cre-mediated recombination in cKO mice.

Since 5-HT_4_R is a G_s_-coupled receptor that activates intracellular cAMP upon ligand binding^10^, we employed a cell culture-based cAMP induction assay to test the signaling ability of the mutant receptor. Expression vectors for mutant *Htr4^delE5^* (delE5) and intact *Htr4* (WT) were made by cloning each transcript from total RNA samples isolated from the hippocampus of a cKO mouse or a WT littermate, respectively. The constructs were transfected into HEK293T cells which were stimulated in the presence of IBMX, a phosphodiesterase inhibitor to allow accumulation of newly converted cAMP. Intact WT expression led to five-fold higher baseline levels of cAMP than of mutant delE5 or EGFP (control) (**Fig. 1d**). Stimulation with zacopride, a potent 5-HT_4_R agonist induced an 88% increase in cAMP production in cells expressing WT but did not significantly alter cAMP levels in those expressing delE5. These results suggest that the deletion of exon 5 leads to a non-functional receptor in vitro.

To test the functionality of mutant 5-HT_4_R in intact hippocampal neurons, we performed whole-cell voltage clamp recordings from dentate gyrus (DG) granule cells (GCs) in acute hippocampal slices from cKO mice and WT littermates. The addition of the 5-HT_4_R agonist, BIMU-8, led to an inward current in WT GCs indicating a depolarizing effect due to the activation of 5-HT_4_R (**Fig. 1e**), consistent with previous reports^11^. In contrast, cKO GCs did not respond to BIMU-8, validating the absence of functional 5-HT_4_R in DG GCs in cKO mice at the electrophysiological level. Together with the cAMP activation assay, these results indicate that the Cre-dependent deletion of exon 5 in cKO mice results in a functionally null receptor.

### Drd3-Cre expression is restricted to mature excitatory neurons in the hippocampus

To determine the identity of the Cre-expressing cells in the hippocampus of Drd3-Cre mice, we crossed them to the Rosa26^fsTRAP^ Cre-dependent reporter line^30^, in which EGFP is fused to ribosomal protein L10a (EGFPL10a). Anti-GFP immunofluorescent staining of coronal brain sections from Drd3-Cre::Rosa26^fsTRAP^ mice revealed labeled cells along the dorsoventral axis of the hippocampus. GFP expression was visible in many cells in the dorsal CA1 (dCA1), dorsal CA3 (dCA3), and ventral CA3 (vCA3) although there were also many unstained cells in these areas (**Fig. 2a**). Few GFP expressing cells were visible in ventral CA1 (vCA1). All of the GFP-expressing cells expressed the neuronal marker, NeuN, but not the GABAergic interneuron marker, glutamate decarboxylase (GAD67; **Fig. S2a**), suggesting Drd3-Cre primarily labels excitatory pyramidal neurons in the CA fields.

**Fig. 2.**
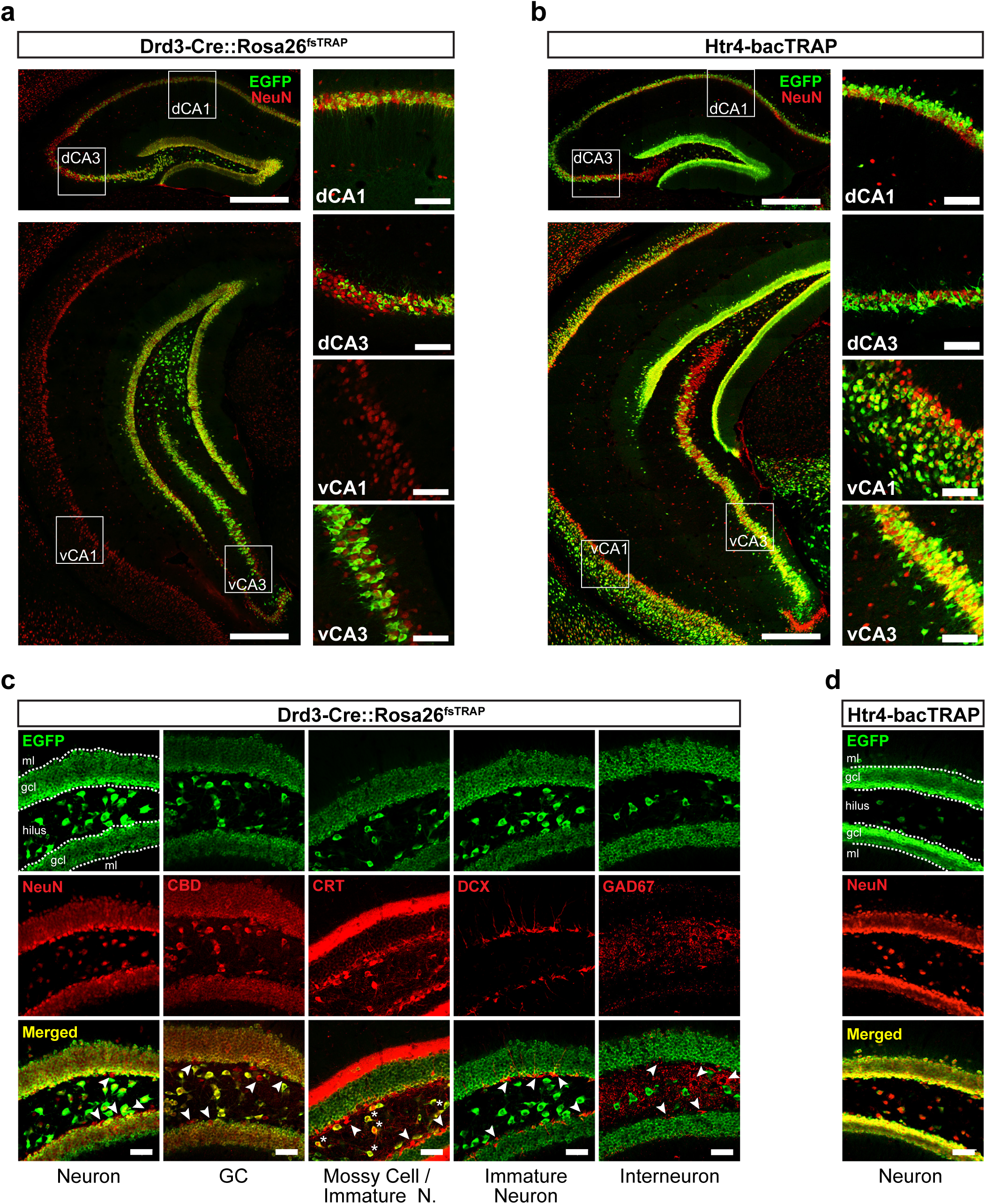
Drd3-Cre expression is restricted to mature excitatory neurons in the hippocampus. **a**, Anti-EGFP (green) and anti-NeuN (red) immunofluorescent confocal images of coronal hippocampal sections at the level of dorsal (top left) and ventral (bottom left) hippocampus from a Drd3-Cre::Rosa^fsTRAP^ mouse. Dorsal and ventral CA1 and CA3 fields (white boxes) are shown at higher magnification on the right. Scale bars: 500 µm (left panels) or 100 µm (right panels). **b**, Anti-EGFP (green) and anti-NeuN (red) immunofluorescent confocal images of coronal sections at the level of dorsal (top left) and ventral (bottom left) hippocampus from an Htr4-bacTRAP mouse. Dorsal and ventral CA1 and CA3 fields (white boxes) are shown at higher magnification on the right. Scale bars: 500 µm (left panels) or 100 µm (right panels). **c**, Immunofluorescent confocal images showing neuronal-type specific marker (red) and EGFP (green) expression in coronal sections through the DG of Drd3-Cre::Rosa^fsTRAP^ mice. Cells labeled with a neuron-type marker but not EGFP (arrowheads) and cells double-labeled with hilar mossy cell marker CRT and EGFP (asterisks) are indicated. ml: molecular layer, gcl: granule cell layer. Scale bars, 50 µm. **d**, Immunofluorescent confocal images of coronal sections through the DG of Htr4-bacTRAP mice labeled with anti-NeuN (red) and anti-EGFP (green). ml: molecular layer, gcl: granule cell layer. Scale bars, 50 µm.

In the DG, GFP expression was visible in the granule cell layer (GCL) and the hilus along the dorsoventral axis (Fig. 2a, c). Co-labeling of GFP with markers for the major DG cell types was used to refine the identity of the Drd3-Cre cells. GFP was detected in all of the calbindin (CBD)-expressing mature GCs and calretinin (CRT)-expressing mossy cells (MCs) in the hilus (asterisks), whereas it was absent from doublecortin (DCX)- and CRT-expressing immature neurons in the GCL and GAD67-expressing GABAergic interneurons throughout the DG (**Fig. 2c**). Together, we conclude that Drd3-Cre targets mature excitatory neurons in the DG, including GCs and hilar mossy cells, but not immature neurons, inhibitory interneurons, or non-neuronal cells.

To determine how the distribution of Drd3-Cre cells compared to that of 5-HT_4_R expressing cells in the hippocampus, we generated Htr4-bacTRAP mice where the EGFPL10a transgene was driven by a *Htr4* BAC promoter to recapitulate endogenous *Htr4* expression (**Fig. S2b**). Like Drd3-Cre mice, GFP+ cells were seen in a subset of cells in the CA fields in the Htr4-bacTRAP mice, although the bacTRAP transgene labeled many more cells in vCA1 (Fig. 2b, d). Also, there was a notable gradient in the number of cells labeled in dorsal and ventral CA3 with the density of labeling decreasing from CA2 to the DG. This pattern can also be seen by ISH (**Fig. S2b** and the Allen Brain Atlas, www.brain-map.org). Interestingly, Drd3-Cre expression shows an opposite gradient with denser labeling near the DG and fewer cells approaching CA2. None of the GFP+ cells co-labeled with GAD67 (**Fig. S2c**). Htr4-bacTRAP expression matched the Drd3-Cre expression in that the vast majority of CBD+ GCs and CRT+ mossy cells co-stained with GFP, but not markers for immature neurons, GABAergic interneurons, progenitor cells, or non-neuronal cells (**Fig. S2d**). Therefore, crossing the Drd3-Cre mice with the Htr4^Floxed^ line results in the loss of functional 5-HT_4_R throughout the DG, but only partial deletion in the CA fields.

### The effect of hippocampus specific loss of 5-HT_4_R on mood and anxiety related behaviors

Given the direct role of serotonin in the regulation of mood^31^ and that 5-HT_4_R has been shown to mediate AD responses^15^, we investigated the impact of the conditional ablation of 5-HT_4_R from hippocampus on mood-related behaviors. We first employed the tail suspension (TST) and forced swim tests (FST) which assess responses to an acute inescapable stressor as a measure of behavioral despair^32,33^. Immobility was decreased in cKO mice compared to WT littermates in both TST and FST (**Fig. 3a, b**), indicating reduced despair-like behavior and mimicking an AD-like response. It is unlikely that these results were due to a general hyperactivity of the cKOs since they did not differ from WTs in locomotor behavior in an open field arena (**Fig. S3e**).

**Fig. 3.**
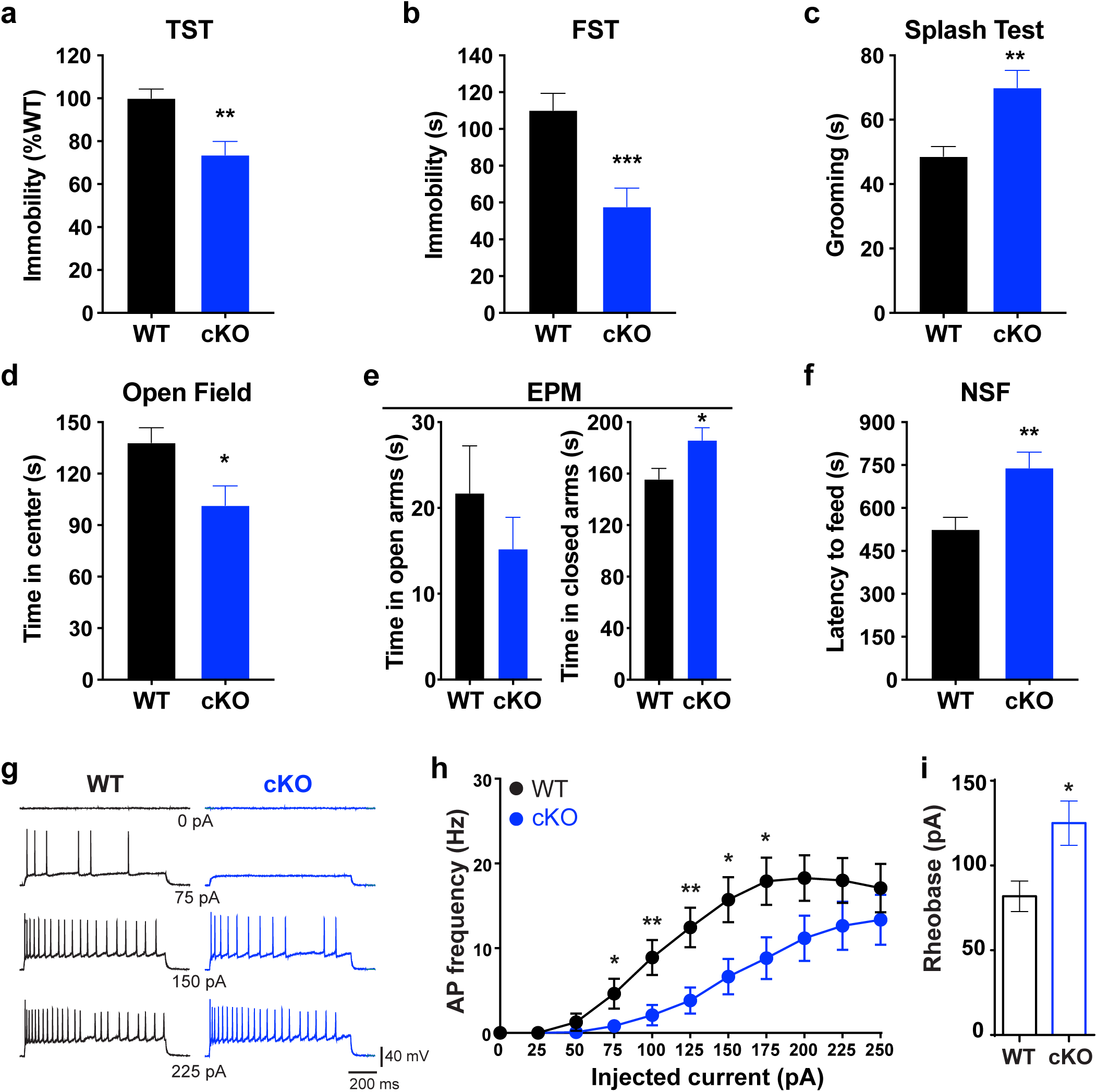
cKO mice exhibit altered affective behaviors and reduced firing in vDG granule cells. **a**, Quantification of the time spent immobile in the tail suspension test (TST) in cKO mice (cKO, blue) and Cre-negative littermates (WT, black). **p = 0.0012, n_WT_ = 19, n_cKO_ = 16. **b**, Quantification of the time spent immobile in the forced swim test (FST) for each genotype. ***p = 0.0005, n_WT_ = 19, n_cKO_ = 16. **c**, Quantification of grooming time in the splash test for each genotype. **p = 0.001, n_WT_ = 19, n_cKO_ = 16. **d**, Mean time spent in the center in the open field (OF) for each genotype. *p = 0.0139, n_WT_ = 19, n_cKO_ = 17. **e**, Quantification of the time spent in the open (left) and closed (right) arms in the elevated plus maze (EPM). *p = 0.0239, n_WT_ = 18, n_cKO_ = 16. **f**, Mean latency to feed in the novelty suppressed feeding (NSF) paradigm for each genotype. **p = 0.0033, n_WT_ = 20, n_cKO_ = 16. **g**, Sample traces from whole-cell current-clamp recordings of GCs in hippocampal slices from WT and cKO mice showing spiking in response to different steps of injected current. **h**, Quantification of the AP frequency of GCs in WT and cKO mice across current steps. **i**, Histogram of rheobase measurements of GCs from WT and cKO mice. All data are represented as mean ± SEM and two-tailed unpaired t-tests were performed for all panels. *p < 0.05, **p < 0.01, ***p < 0.001.

The splash test and sucrose consumption are commonly used as indicators of hedonic state in depression models and rely on the animal’s motivation for self-grooming and the rewarding properties of a sweet, palatable sucrose solution, respectively^34–36^. In the splash test, cKO mice spent more time grooming following the application of a sucrose solution compared to WT littermates, mimicking the effect of antidepressant administration (**Fig. 3c**). During the sucrose consumption, both cKO and WT mice drank approximately 5-10-fold more sucrose than water each day over the three-day test (**Fig. S3a, b**), implying no difference in hedonic capacity between genotypes.

Interestingly, cKO mice consumed cumulatively more sucrose after 72 hours compared to WT. This result was unlikely due to a change in caloric need since body weight was unaltered in the cKOs (**Fig. S3c**). The significance of the increased sucrose consumption is difficult to interpret given the strong sucrose preference exhibited by both groups but may reflect a subtle rise in pleasure-seeking drive. Together with the FST and TST results, these data indicate that cKO mice displayed AD-like behaviors in despair- and hedonic-based assays.

As anxiety is closely associated with mood disorders and AD treatment, and has been linked to hippocampal and 5-HT_4_R function^18,37,38^, we next assessed how hippocampus-specific loss of 5-HT_4_R affected a variety of anxiety-related behaviors. The open field test (OF), elevated plus maze (EPM) and novelty suppressed feeding (NSF) paradigms all rely on mice having an innate aversion to well-lit, open spaces while NSF adds the conflict of engaging in feeding behavior in a novel environment^39–41^. In the OF, cKO mice spent less time the center of the arena during the initial 10 min compared to WT littermates while showing similar overall locomotor activity (**Fig. 3d, Fig. S3d, e**). Similarly, in the EPM, the cKOs spent more time in the closed arms compared to WT (**Fig. 3e**). Although the time spent in the open arms did not differ between genotypes, the cKOs displayed a trend towards higher velocity in EPM compared to WT (**Fig. S3f**) suggesting that higher exploratory activity of cKO mice was not accompanied by more exploration of the open arms. The cKO mice also displayed a greater latency to feed in NSF (**Fig. 3f**) yet consumed more food on average when placed back in their familiar home cages following NSF (**Fig. S3g**), conveying that the lack of motivation to feed was not the source of their prolonged latency. Together, these conflict-based behavioral analyses indicate that cKO mice have an anxiety-like phenotype.

We also tested behaviors that may be affected by anxiety levels such as social interaction and acoustic startle response (ASR). Mice were tested in the three-chamber social interaction test to measure sociability since a reduced amount of time spent interacting with a novel conspecific has been associated with abnormal anxiety-like behaviors^42^. We observed that cKO mice interacted less with a novel mouse compared to WT (**Fig. S3h**). Enhanced startle response has also been linked to increased anxiety and its physiological features^43,44^ while blunted startle was observed in patients with anhedonia^45–47^. The acoustic startle reflex, a defensive motor response to an intense sudden auditory stimulus, is measured to assess ASR in mice. We found that ASR was significantly enhanced in cKO mice compared to WT (**Fig. S3i, j**). On the other hand, the pre-pulse inhibition (PPI), the suppression of ASR when a startling stimulus is preceded by an immediate, weaker pre-stimulus, was not altered (**Fig. S3k**). Together, these anxiety-regulated responses reflect the anxiogenic phenotype observed in the cKOs.

### Loss of 5-HT_4_R reduces the excitability of dentate gyrus granule cells

Activation of 5-HT_4_R leads to depolarizing currents via second messenger mechanisms in excitatory cells of the hippocampus^11^, including DG GCs (**Fig.1d**). To test if the cell type specific loss of functional 5-HT_4_R had any electrophysiological consequences, we performed whole-cell current clamp recordings from vDG GCs in acute hippocampal slices from WT and cKO mice. We found that the firing frequency of GCs was significantly reduced in cKO slices compared to WT (**Fig. 3g, h**). In addition, rheobase, the amount of current necessary to be injected to elicit an action potential, was larger in GCs from cKO mice (**Fig. 3i**). On the contrary, there was no difference in the properties of single action potentials between genotypes, including threshold, amplitude, half-amplitude width and fast after-hyperpolarization as well as the input resistance and slow after-hyperpolarization (**Fig. S4a-g**). These data show that the loss of 5-HT_4_R from DG GCs leads to specific physiological changes resulting in a reduced excitability in these neurons.

### Molecular adaptations in the vDG of cKO mice highlight cellular and functional roles for 5-HT_4_R

We next investigated global differences in gene expression that occurred following the deletion of hippocampal 5-HT_4_R to identify molecular mechanisms that may underlie the behavioral and electrophysiological phenotypes observed in cKO mice. We focused on the ventral DG since nearly all 5-HT_4_R expressing cells in the DG, but not CA fields, are targeted by the Drd3-Cre line (**Fig. 2a-c**), and there is a strong link between ventral hippocampus and the regulation of depression and anxiety related behaviors^27^. The translating ribosome affinity purification (TRAP) method^48,49^ was used to isolate cell type specific transcripts by expressing the EGFPL10a TRAP transgene in WT and cKO Drd3-Cre cells by crossing Drd3-Cre with Rosa26^fsTRAP^, and Drd3-Cre::Htr4^loxP/loxP^ with Rosa26^fsTRAP^::Htr4^loxP/loxP^ mice, respectively (**Fig. 4a**). Polysome-bound mRNAs were extracted following anti-EGFP immunoprecipitations (IP) on vDG homogenates and analyzed by RNA-seq alongside mRNAs extracted from whole tissue (input) (**Fig. S5a**). The quality of dissection was confirmed by qRT-PCR which showed that markers for DG (*Dsp)* and vDG (*Trhr*) were highly expressed in vDG inputs but markers for CA fields (*Tyro*) and dDG (*Lct*) were not (**Fig. S5b-c**). Visualization of RNA-seq reads mapped to the *Htr4* locus confirmed the deletion of exon 5 in cKO tissue (**Fig. 4b**), although overall expression of *Htr4* was increased in cKO TRAP mRNA compared to WT, likely reflecting an attempt by the cells to compensate for the non-functional protein (**Fig. 4c**). There did not appear to be a broader compensation in 5-HT signaling since there was no difference in the expression of other 5-HT receptor transcripts between genotypes (**Fig. 4c**). Consistent with previous reports of the distribution of 5-HT receptors in the DG^28^, the receptors with highest expression in the vDG TRAP were *Htr1a*, *Htr2a*, *Htr4*, and *Htr5a*.

**Fig. 4.**
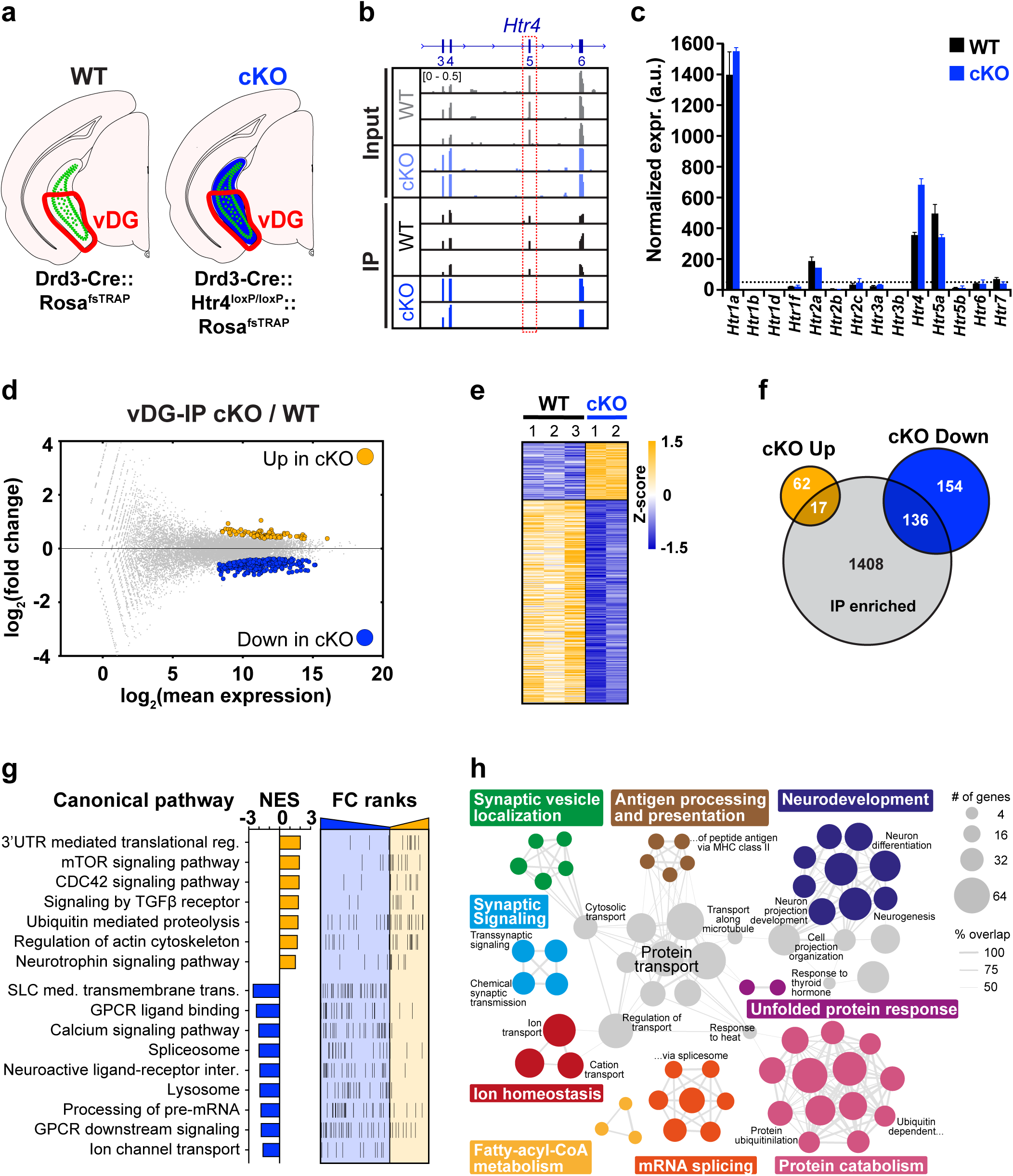
TRAP profiling of vDG neurons in the absence of 5-HT_4_R. **a**, Schematic depicting the cell types targeted for TRAP. Green represents EGFPL10a expressing cells and blue indicates conditional 5-HT_4_R deletion. The area dissected is outlined in red. **b**, Browser view of mapped reads showing the lack of expression of *Htr4* exon 5 in IP and input samples in the cKO. **c**, Bar graph of normalized expression values (mean ± SEM) of all 5-HT receptors (Htrs) in IP samples. Dashed line is drawn at normalized expr. = 50. Only Htr4 has FDR < 0.1. **d**, MA-plot highlighting differentially expressed (DE) genes (FDR < 0.1) between the cKO and WT TRAP mRNA. Complete list of IP DE genes can be found in Table S1. **e**, Heatmap visualization of DE genes between genotypes. **f,** Venn diagram showing the overlap of up- and down-regulated DE genes from **e** with genes enriched in the Drd3-Cre-expressing cells (IP enriched, Table S2). **g**, Summary of gene set enrichment analysis (GSEA) performed on the fold change (FC)-ranked gene list (p<0.05) between cKO and WT IP data sets. The normalized enrichment score (NES) shows the direction of enrichment in the cKO and FC-ranks are arranged from down-regulated (blue) at left to up-regulated (yellow) at right. Complete list of pathways can be found in Table S3. **h**, Summary of gene ontology (GO) analysis of DE from **e**. All categories shown have FDR < 0.01, full analysis is provided in Table S4.

A direct comparison of gene expression between TRAP mRNA (IP) and whole tissue mRNA (input) revealed a substantial depletion of markers for interneurons, neurogenesis, and other non-neuronal cells in the TRAP data set (**Fig. S5d, e**), consistent with our immunohistology showing the Drd3-Cre line primarily labels mature GCs and hilar MCs in the DG (**Fig. 2c**). While markers for GCs were highly expressed in TRAP mRNA, their levels of enrichment in IP over input were less robust compared to the depletions of markers of cells not labeled by Drd3-Cre (**Fig. S5e, f**), likely because mature GCs constitute the vast majority of all cells in the DG. A similar distribution of marker genes was observed for the cKOs (**Fig. S5d**) and genes enriched or depleted in IP compared to input were similar between WT and cKO (**Fig. S6a**) indicating that the mutant allele did not noticeably alter cellular identity.

Differential expression analysis of TRAP mRNA between WT and cKO mice identified 80 upregulated and 290 downregulated genes following the loss of 5-HT_4_R function (**Fig. 4d**). Heatmap visualization based on normalized expression values showed that the significant gene expression differences between genotypes were consistent across all replicates (**Fig. 4e**). Over 40% of the differentially expressed (DE) genes (17 up- and 136 downregulated) were enriched in the IP of Drd3-Cre cells (**Fig. 4f**), illustrating a strong cell type specific adaptation. To gain a general understanding of the gene pathways dysregulated in the cKOs, we performed gene set enrichment analysis (GSEA) on a ranked list of genes differing between WT and cKO samples (**Fig. 4g**). We found positive enrichment in signaling pathways linked to translation, neuroplasticity and cytoskeletal regulation, including mTOR, CDC42, TGFβ, neurotrophin signaling and regulation of actin cytoskeleton, all of which have been shown to be regulated by AD treatment as well. Interestingly, there was an upregulation in ubiquitin proteolysis and 3’UTR mediated translation, and a downregulation in pre-mRNA processing, implying a possible response to the handling of mutant *Htr4* transcript and 5-HT_4_R protein. We also found negative enrichment in pathways related to 5-HT_4_R function including GPCR ligand binding, calcium signaling, neuroactive ligand receptor interaction, and GPCR downstream signaling, paralleled by decreased expression of genes involved in cAMP signaling (**Fig. 4g, S6b**). In addition, there was a downregulation in pre-mRNA processing and transmembrane transport of various classes of ions (**Fig 4g, S6c**). Functional annotation analysis of the list of genes that were significantly differentially expressed between genotypes further showed alterations in a variety of processes underlying neuronal function including synaptic vesicle localization, neuron development, synaptic signaling, ion homeostasis, and metabolism while unfolded protein response, protein transport, and protein catabolism imply a change in proteostasis (**Fig. 4h**). Together, our differential expression analysis of the vDG from cKO and WT mice identified substantial transcriptional changes in the absence of 5-HT_4_R that are related to neuronal function and intracellular 5-HT_4_R receptor signaling.

### The ablation of 5-HT_4_R resulted in enhanced adult hippocampal neurogenesis specifically in the vDG

Up to this point, we have investigated the effects of 5-HT_4_R loss on the intrinsic molecular and electrophysiological properties *Htr4*-expressing cells. We next asked how these adaptations impacted the overall cellular environment within the DG. To do so we took advantage of the whole tissue mRNA (input) since it was enriched with transcripts expressed in non-neuronal cells, progenitors, and interneurons (**Fig. S5e**) allowing us to examine broad molecular responses in non-*Htr4*-expressing cells. Differential expression analysis between WT and cKO tissue revealed a robust molecular response (625 DE genes) in the whole vDG and a relatively small response in the whole dDG (179 genes) (**Fig. 5a, S7a**). Only 25 genes were shared between regions (**Fig. S7b, c**). Functional annotation of DE genes revealed remarkably distinct cellular adaptations between the two regions. Processes related to neural function and plasticity, including neurogenesis, neuron development and differentiation, synaptic signaling, ion transport and homeostasis, and regulation of action potential where uniquely enriched in the whole vDG mRNA but not dDG (**Fig. 5b, S7d**). The genes altered in the whole dDG mRNA were primarily related to protein and RNA processing. Furthermore, disease-gene association analysis of DE genes in the whole vDG also showed significant enrichment for a variety of diseases, including affective disorders such as depressive, bipolar and anxiety disorders, along with other psychiatric disorders (**Fig. S7e**). No significant disease-gene associations were found in the dDG.

**Fig. 5.**
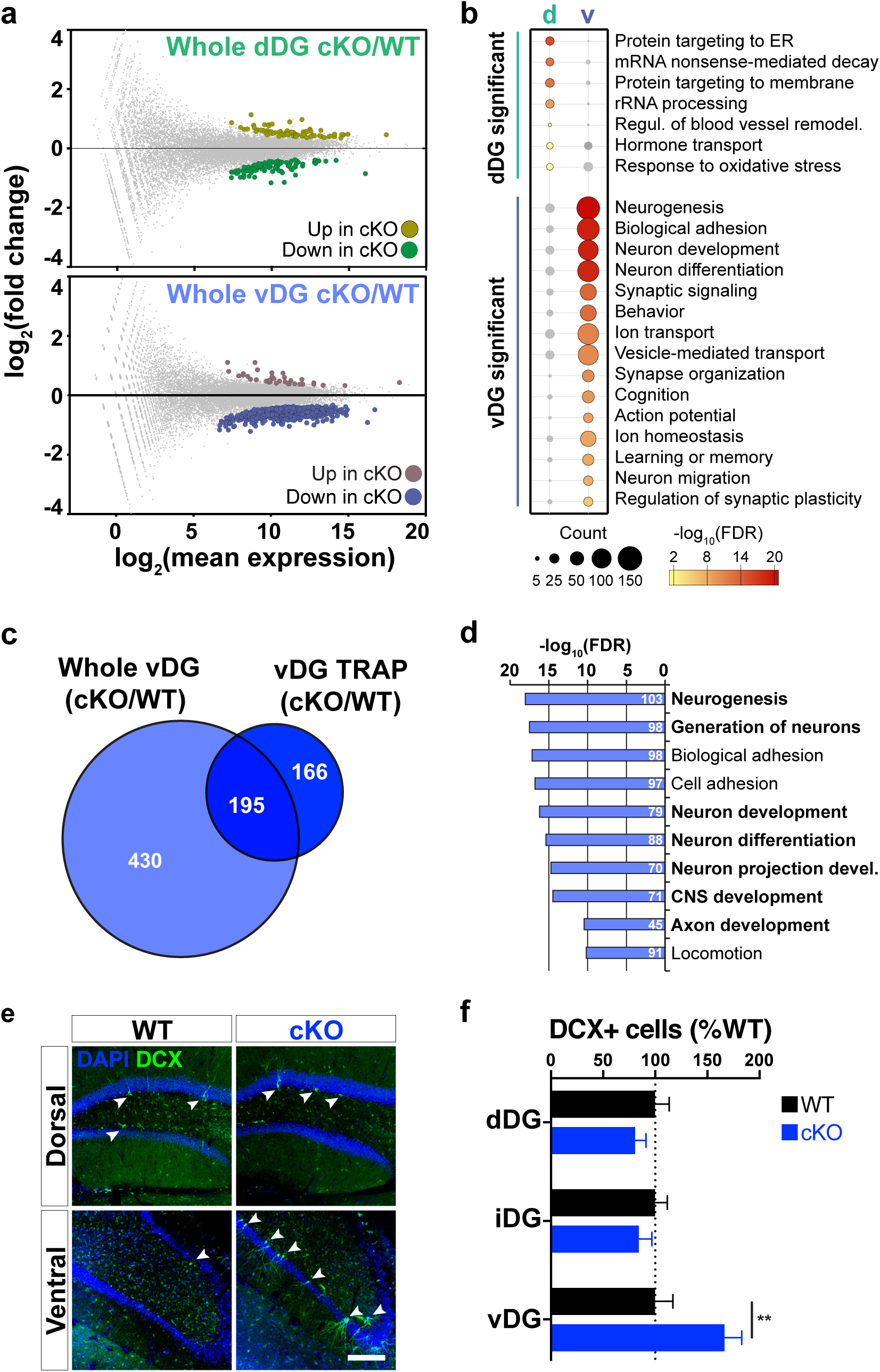
Loss of 5-HT_4_R led to changes in neurogenesis-related genes and increased number of immature neurons in the vDG. **a**, MA-plots indicating differentially expressed (DE) genes (FDR < 0.05) between cKO and WT mRNA from whole dDG (top) and vDG (bottom). Complete lists can be found in Table S5 and S6. **b**, Comparison of significantly enriched gene ontology (GO) terms among whole dDG and vDG DE genes from **a**. **c**, Venn diagram comparing DE transcripts shared and unique to vDG TRAP (dark blue) and whole vDG tissue (light blue) samples. **d**, Top ten GO terms enriched in the whole vDG-specific gene list from **c**. Terms related to neurogenic tone are in bold. **e**, Representative images of anti-DCX immunofluorescence in coronal sections through dDG and vDG from cKO and WT mice. Arrowheads indicate DCX+ cells. DAPI is shown in blue. Scale bar, 150 µm. **f**, Quantification (mean ± SEM) of DCX+ cells per granule cell layer area in the dorsal (dDG), intermediate (iDG), and ventral (vDG) DG. Two-way ANOVA: genotype × region interaction, F(2,24) = 6.328, p = 0.0062 followed by post hoc Fisher’s LSD test, **p = 0.002. n = 5 per genotype.

To identify the gene expression changes that occurred primarily in non-*Htr4*-expressing cells in the cKOs, we directly compared the DE genes from the vDG TRAP (IP) and whole vDG mRNA (input), and found 430 genes that were altered in the whole vDG but not TRAP (**Fig. 5c**). These whole vDG-specific genes were predominantly related to neurogenesis and neurodevelopment, likely reflecting the fact that progenitors and immature neurons comprised many non-Cre-labeled cells (**Fig. 5d**). It should also be noted that many of the cell intrinsic processes, such as ion homeostasis and transport, synaptic signaling, and protein processing appeared to be unique to the TRAP (**Fig. 4e, f and 5b**).

Our genome-wide transcriptional analysis revealed significant alterations in gene expression related to neurogenesis in the whole vDG and an upregulation of factors known to promote neurogenesis and plasticity specifically in the Drd3-Cre cells (**Fig. 4e, f** and **5b, d**). This is particularly interesting given that the cKOs exhibit AD-like behavioral responses and the link between adult neurogenesis of DG granule cells and antidepressants^27^. We therefore examined whether we could detect any change in hippocampal neurogenesis in the cKOs. The number of doublecortin-positive (DCX+) immature neurons along the dorsoventral axis of the subgranular zone (SGZ) and GC layer of the DG was quantified in WT and cKOs (**Figs. 5e** and **S7f**). There was a 67% increase in the number of DCX+ neurons in the vDG of cKO mice compared to WT littermates, while no difference in the number of DCX+ neurons was detected in the dDG or intermediate DG (**Fig. 5f**). These results suggest that loss of functional 5-HT_4_R led to enhanced neurogenesis in the vDG, which likely occurred through a non-cell-autonomous manner since Drd3-Cre does not target progenitor cells or immature neurons. Together, these data demonstrate a remarkable amount of region-specific cellular and molecular adaptations along the dorsoventral axis of DG in response to loss of 5-HT_4_R function.

## Discussion

In the current study, we generated conditional 5-HT_4_R knockout mice to examine the role of this receptor in discrete cell populations in the hippocampal formation at behavioral, cellular, and molecular levels. The loss of 5-HT_4_R primarily from mature excitatory neurons in the DG and hilus led to AD-like performance in behavioral tests of despair and anhedonia, and elevated anxiety. DG granule cells were less excitable and showed altered expression of genes related to GPCR function, ion transport, and neuroplasticity in the absence of 5-HT_4_R. In addition, cKO mice showed increased neurogenesis in the vDG but not dDG, indicating regional differences in adaptions to 5-HT_4_R loss in the local environment along the dorsoventral axis. Taken together, these results strongly suggest that the removal of functional 5-HT_4_R from mature excitatory cells in the DG mimics chronic AD treatment and offers insight into hippocampal circuits modulating mood and anxiety.

Decreased functionality of hippocampal 5-HT_4_ receptors is consistent with reports describing cellular responses to antidepressants. Chronic SSRI treatment has been shown to reduce the density of 5-HT_4_R binding and decrease 5-HT_4_R-dependent cAMP signaling and postsynaptic excitability in hippocampal pyramidal cells in rats^50^. Moreover, short-term treatment with the partial 5-HT_4_R agonist, RS67333, leads to the desensitization of the receptors in hippocampus^17^. These findings together with our results paint the picture of a multifaceted role for 5-HT_4_R in the regulation of neural circuits underlying mood. 5-HT_4_R is expressed in many limbic structures besides the hippocampus including prefrontal cortex, ventral striatum, and amygdala^16^. Indeed, the mPFC has been implicated as a site mediating the fast-acting anxiolytic/antidepressant effects of RS67333^19,51^ and the overexpression of 5-HT_4_R specifically in the mPFC led to AD-like behavior^52^. In addition, we previously found that chronic SSRI administration led to an induction of 5-HT_4_R expression in corticostriatal neurons in sensorimotor cortex^5^, although it does not appear to underlie SSRI-mediated changes in 5-HT responses in those cells^53^. The lack of impact of constitutive deletion of 5-HT_4_R on baseline mood^21^ further attests to the complex interplay between different circuit components that express the receptor.

Our results showed that the absence of 5-HT_4_R led to a reduced electrophysiological excitability of DG GCs. It is not clear whether the changes we observed in the cKOs were due to a deficiency of 5-HT_4_R signaling directly or a result of long-term cellular adaptations in the absence of the receptor. 5-HT_4_R has been shown to contribute to the excitability of pyramidal cells through PKA-mediated closure of Ca^2+^-activated K^+^ channels underlying sAHP current^11,54–56^, however we did not see any change in slow or fast AHPs in the cKOs. A direct coupling to both voltage-sensitive calcium channels^57^ and the extracellular signal-regulated kinase pathway^58^ has also been proposed for 5-HT_4_R function, both of which are known to modulate neuronal function^59,60^. The TRAP data revealed a dysregulation of Ca^2+^ signaling, ion homeostasis, and/or ion channel expression in the 5-HT_4_R cKOs, although more work is needed to understand the significance of these changes on the excitability of the GCs.

The AD-like phenotype we observed following the deletion of 5-HT_4_R in the hippocampus is possibly mediated by the decreased excitability of the vDG GCs observed in the cKOs. Growing evidence supports a link between suppressed activity of DG GCs and AD-like behavior. Decreased excitability of vDG GCs through local GABAergic inhibition produces behavioral AD action^61^, and indirect inhibition of mature GCs of the vDG by young adult-born GCs (abGCs), which are increased by AD treatment, or by chemogenetic methods promotes resilience to stress^62^. In addition, SSRI-mediated inhibition of mature GCs through 5-HT_1A_R is necessary for AD responses in preclinical models^63^. Blockade of GC-CA3 glutamate release and NMDA receptor activity reduces behavioral despair^64^, and the therapeutic effects of antidepressants and 5-HT_4_R agonists in rodents are dependent on the tonic inhibition of CA3 pyramidal cells via the activation of 5-HT_1A_ receptors^15^. Finally, chronic AD administration attenuates excitatory physiological effects of 5-HT_4_R throughout the hippocampal circuit^50,65^. Although we did not detect a compensatory change in the expression of 5-HT_1A_ or other 5-HT receptor transcripts in GCs following 5-HT_4_R deletion, it is possible that the AD-like phenotype of the cKOs resulted from a shift in the balance of excitatory and inhibitory 5-HT signaling. 5-HT_4_ and 5-HT_1A_ receptors are both highly expressed in DG GCs and clearly cooperate to mediate cellular responses to 5-HT despite activating opposing second messenger pathways and electrophysiological properties^9,63^. The broad downregulation of cAMP-related genes and GPCR signaling detected in the TRAP data supports this idea.

Behavioral analysis of the cKOs also reinforced a central role for hippocampal 5-HT_4_R in modulating anxiety. Previous studies using systemic administration of pharmacological agents demonstrated robust anxiolytic properties of 5-HT_4_R agonists while 5-HT_4_R antagonists blocked the anxiolytic effects of diazepam and SSRIs^16,18,66^. Although there is a discrepancy in early reports also showing anxiolytic effects of 5-HT_4_R antagonists in rodents^66,67^, no obvious anxiety-like phenotype was observed in constitutive 5-HT_4_R KO mice^21^. The anxiogenic phenotype observed in our 5-HT_4_R cKO mice implies hippocampal 5-HT_4_R directly moderates anxiety, possibly though its role in modulating the excitability of vDG GCs. Acute optogenetic activation of vDG GCs was shown to suppress innate anxiety, although increased anxiety was not observed when vDG GCs were acutely inhibited^38^. Long-term inhibition of GCs due to chronic loss of 5-HT_4_R functionality in the cKOs may have amplified this effect in our study. The activity of discrete subsets of vCA1 projection neurons have differing effects on anxiety-related behaviors depending on their postsynaptic targets^68,69^, indicating a complexity of cell specific mechanisms even within the hippocampal circuit. Although Drd3-Cre was absent from vCA1 and 5-HT_4_R function was likely preserved in most vCA1 pyramidal cells in the cKOs, reduced activity of vDG GCs has a significant impact on the overall trisynaptic hippocampal circuit and the outputs of vCA1 pyramidal cells.

5-HT_4_R deletion also had a palpable effect on molecular and cellular neuroplasticity in the vDG. Cell type specific molecular profiling of vDG GCs with TRAP was able to detect a clear downregulation of GPCR signaling and an increase in protein degradation pathways, reflecting direct intrinsic adaptations to the loss of functional 5-HT_4_R and processing of the unstable mutant protein. However, it also provided evidence for an alteration in the neuroadaptive state of the cells highlighted by a change in genes linked to processes such as neuronal development, differentiation, and synaptic signaling. These pathways have been shown to be differentially regulated in the hippocampus with AD treatment, deep brain stimulation, and exposure to stress^7,8, 70–73^ and likely underlie the changes in synaptic plasticity, spine density, and dendritic architecture of hippocampal neurons observed after AD administration^3,74,75^.

The molecular adaptations in mature GCs were accompanied by heightened generation of abGCs selectively in the vDG in the cKOs. Hippocampal neurogenesis is a reliable readout of AD activity^22^ and mediates some of the behavioral effects of AD treatments and responses to stress^35,62,76^ but does not produce AD-like behavior on its own^77^. The absence of 5-HT_4_R and Cre from neural progenitors and immature GCs suggests the enhanced neurogenesis seen in the cKOs was due to non-cell autonomous mechanisms, likely resulting from a pro-neurogenic change in the local environment within the vDG. This notion is strongly supported by the enrichment of neurogenesis-related pathways being highly regulated in the vDG niche of the cKOs. The lack of change in neurogenesis at the molecular and cellular level in the dDG underscores the functional divergence of the DG along the dorsoventral axis^27,78^ and the regional specificity of 5-HT_4_R function. A similar anatomical discrepancy in neurogenesis was observed in patients and mice following chronic SSRI treatment^79,80^ and the ablation of abGCs in the vDG, but not dDG attenuated some of the behavioral responses to AD treatment, stress, and chronic pain^81,82^.

The present study begins to refine the role of 5-HT_4_R in a key circuit that mediates mood and anxiety and reinforces the complexity of neuromodulatory signaling in arbitrating emotion. This notion is underscored by the fact that the AD-like phenotype of the conditional deletion of 5-HT_4_R from mature granule cells in the DG is accompanied by an increase in anxiety. SSRIs and other AD drugs are commonly used to treat anxiety disorders so the anxiogenic behavior of the 5-HT_4_R cKO mice seems to oppose their AD-like phenotype, indicating a dissociation between the mechanism underlying each process. Indeed, antidepressants can promote suicidality, anxiety and nervousness when given to healthy subjects^83^, and acute and chronic SSRI treatment in rodents can induce anxiety-like behaviors depending on the baseline anxiety/stress/depression levels, strain, age, dose and delivery method of AD treatment^84–87^. Our data justify further inquiry to determine if these phenomena are moderated by a disengagement of hippocampal 5-HT_4_R. Such observations and the widespread influence of the therapeutic of targeting neuromodulatory systems underscore the need to tease apart regional and cell type specific mechanisms to develop better drugs with diminished side effects.

## Acknowledgements

This work was funded by NIH/NINDS grants R01NS091722 (E.F.S.) and R21NS105047 (E.F.S.), NIH/NIDA P30 Center DA035756 (N.H., E.F.S.), The Fisher Center for Alzheimer’s Research Foundation (P.G.) and The Leon Black Family Foundation (P.G.). N.H. is also an investigator of the Howard Hughes Medical Institute (HHMI). We thank The Rockefeller University Genomics Resource Center, Transgenic Services Laboratory and Comparative Bioscience Center as well as the HHMI Janelia Research Campus Gene Targeting and Transgenics support team. We also thank Erika Andrade and Awni Mousa for assistance with bioinformatics analyses and Jodi Gresack for assistance and discussions regarding the behavioral studies. We thank Winrich Friewald for guidance and David Rockefeller Graduate Program for their support.

## Author contributions

R.K., N.H., and E.F.S designed the study. R.K. performed all experiments unless noted otherwise. L.M. conducted electrophysiology experiments. M.N.R performed FISH experiments. E.B.H assisted with behavior and RNA-seq. R.K., L.M., J.L.W-S, P.G., N.H., and E.F.S. analyzed data. R.K., L.M., and E.F.S. wrote the manuscript.

---

**Fig. S1.**
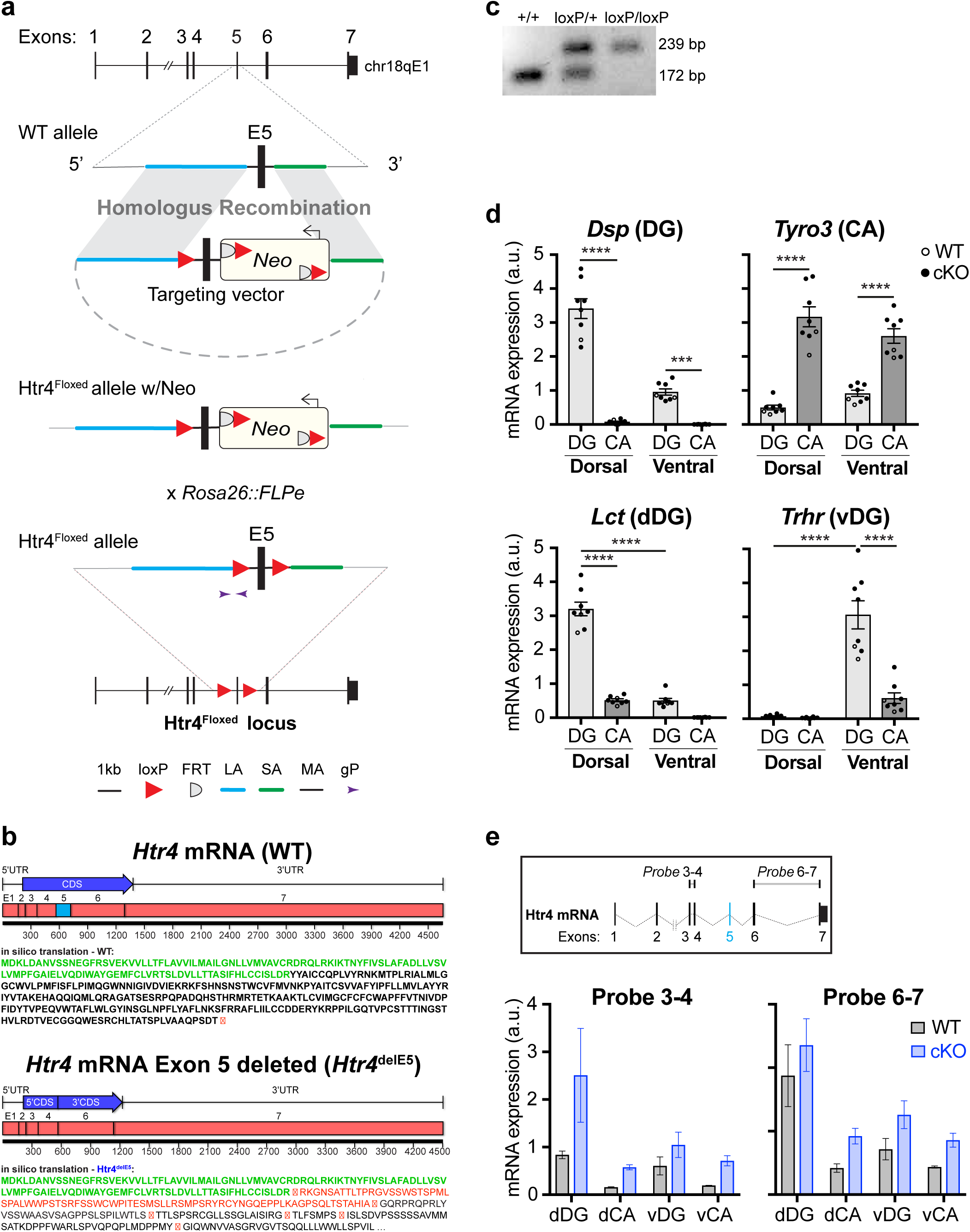
Gene targeting strategy to generate Htr4^Floxed^ mouse line. Related to Fig. 1. **a**, Schematic of *Htr4* gene targeting steps. Targeting vector was designed to drive the homologous recombination via long (LA, blue) and short (SA, green) arms to insert the loxP-flanked exon 5 (vertical black bar) and Neomycin resistance cassette (Neo, yellow rectangle). Relative positions of FRT (flippase recognition target) sites (grey half circle) and loxP sites (red triangle) are shown. FLPe-mediated recombination was used to excise the Neo cassette and lead to *Htr4* exon 5 flanked with loxP sites (Htr4^floxed^ allele). gP, genotyping primers (purple arrowheads). Scale bar for magnified schematics, 1kb. **b**, In silico translation analysis predicting that the loss of exon 5 from *Htr4* mRNA would lead to immediate early stop codons. Green amino acid sequence represents the translation up to exon 5. Red squared X represents the translation of the stop codons. **c**, Sample genotyping results using gP from a for mice that are Htr4^+/+^ (+/+), Htr4^+/loxP^ (loxP/+) and Htr4^loxP/loxP^ (loxP/loxP) showing the shift in amplicon size in the targeted allele. **d**, Quantitative RT-PCR (mean ± SEM) of the expression of marker genes for each hippocampal region in all samples used in Fig. 1c. Data points represent biological replicates of WT (empty circles, n = 2) and cKO (filled circles, n = 6) samples. One-way ANOVA followed by post hoc Fisher’s LSD test, ***p < 0.001, ****p < 0.0001. **e**, Top shows a schematic of TaqMan probes spanning *Htr4* exons 3-4 (Probe 3-4), and 6-7 (Probe 6-7). Bottom is quantitative RT-PCR results (mean ± SEM) showing the persistent expression of exon 3, 4, 6 and 7 in *Htr4* mRNA in cKO mice compared to WT along the dorsoventral axis of the hippocampus.

**Fig. S2.**
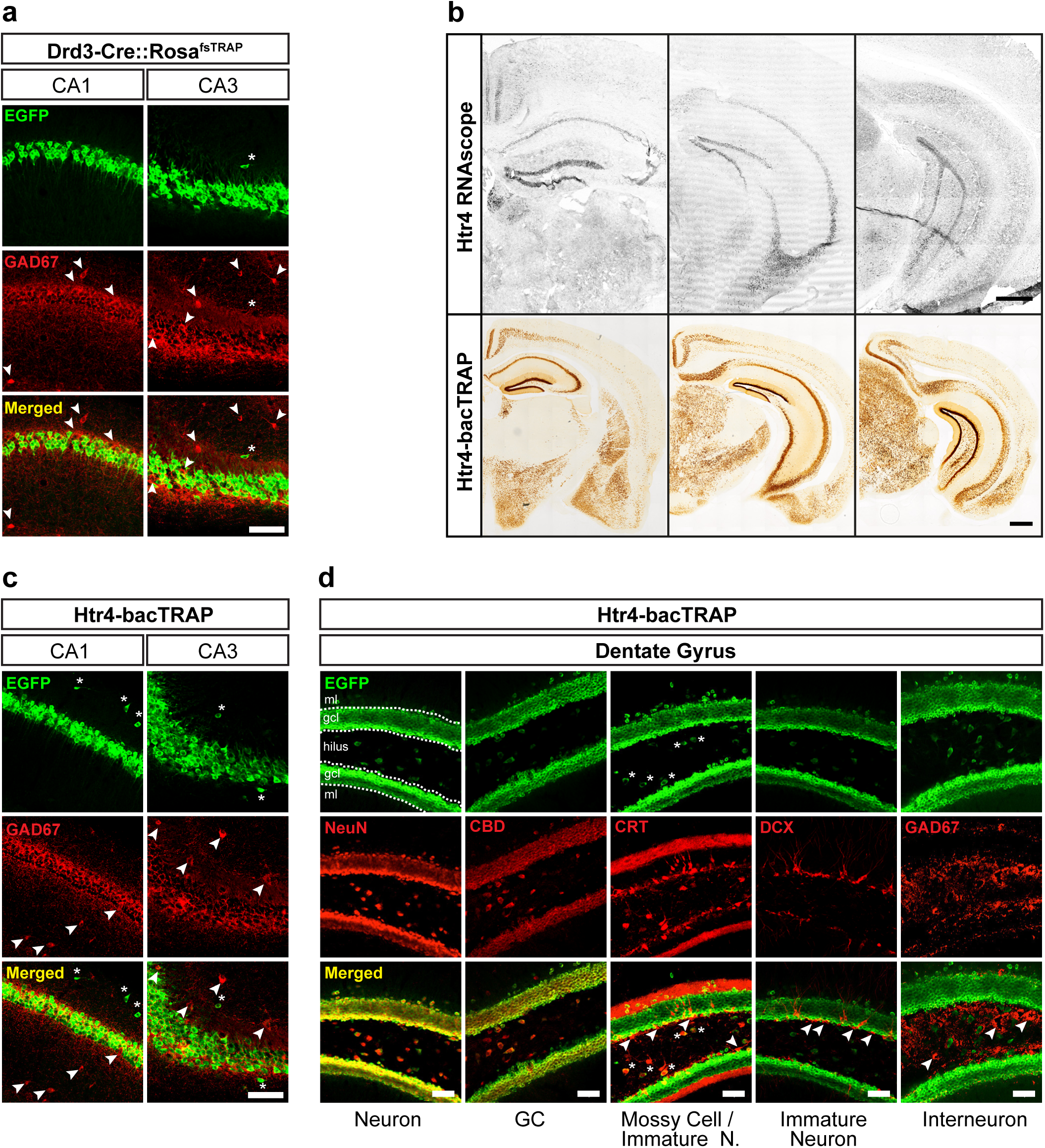
Drd3-Cre and Htr4-bacTRAP expression is restricted to mature excitatory neurons in the hippocampus. Related to Fig. 2. **a**, Anti-EGFP (green) and anti-GAD67 (red) immunofluorescent confocal images of coronal hippocampal sections from a Drd3-Cre::Rosa^fsTRAP^ mouse. Drd3-Cre driven EGFP expressing cells were GAD67-negative, both in the CA1 (left), and CA3 (right) fields. Arrowheads, GAD67-postive; asterisks, GAD67-negative. Scale bar, 100 µm. **b**, Above, FISH images showing the spatial expression profile of *Htr4* in the mouse brain. Below, Anti-EGFP DAB immunohistochemistry of a Htr4-bacTRAP mouse brain showing that the expression of the BAC transgene EGFPL10a under the regulatory elements of *Htr4* recapitulate the expression profile of *Htr4* in the brain. **c**, Anti-EGFP (green) and anti-GAD67 (red) immunofluorescent confocal images of coronal hippocampal sections from a Htr4-bacTRAP mouse. EGFP expressing cells were GAD67-negative, both in the CA1 (left), and CA3 (right) fields. Arrowheads, GAD67-postive; asterisks, GAD67-negative. Scale bar, 100 µm. **d**, Immunofluorescent confocal images showing neuronal-type specific markers (red) and EGFP (green) expression in coronal DG sections of Htr4-bacTRAP mice. Arrowheads, cells expressing a specific neuron-type marker but not EGFP. Asterisks, cells coexpressing hilar mossy cell marker CRT and EGFP. ml: molecular layer, gcl: granule cell layer. Scale bars, 50 µm.

**Fig. S3.**
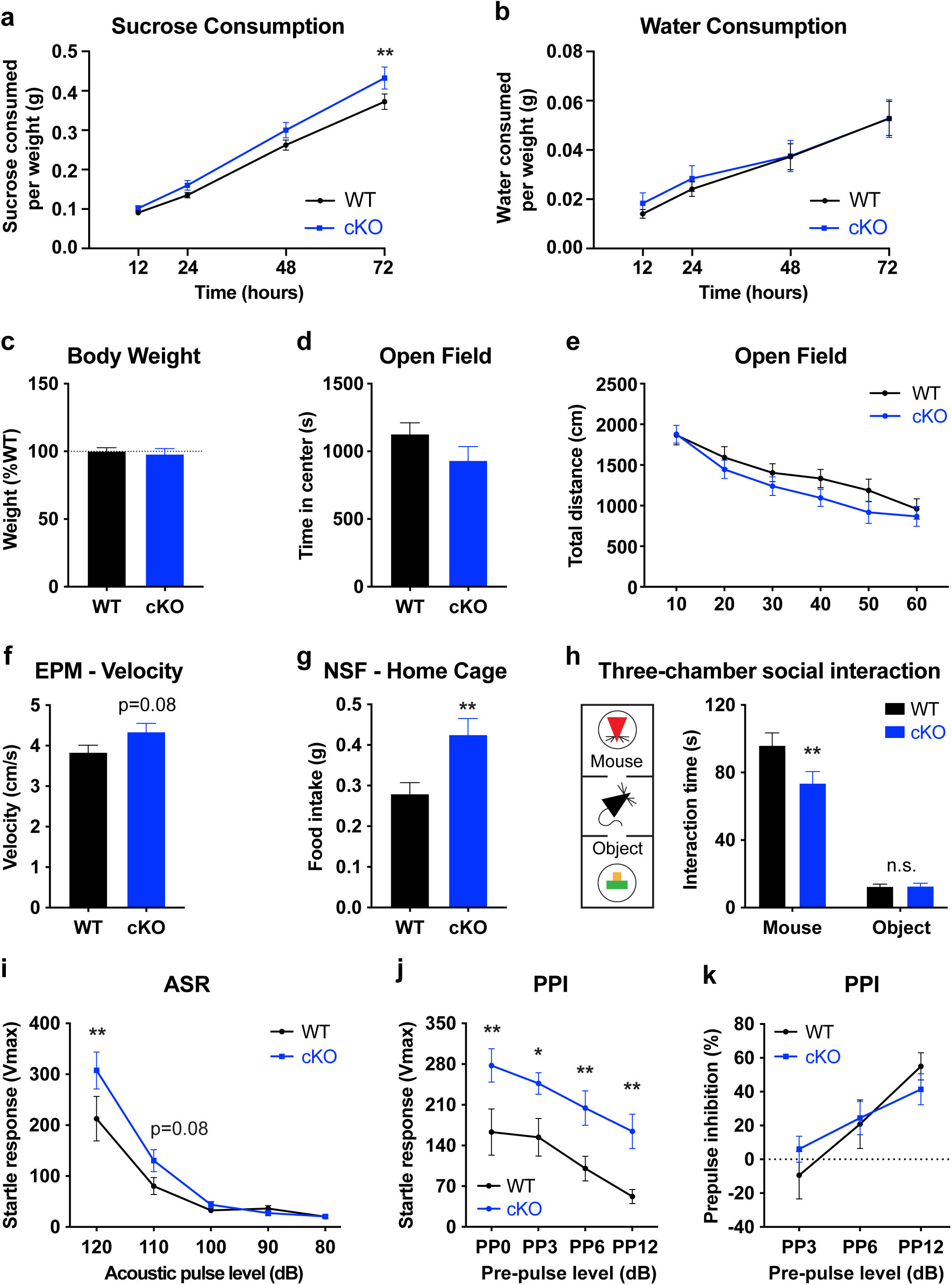
Additional behavioral assessment of cKO mice. Related to Fig. 3. **a**, Quantification of the amount of 1% sucrose solution consumed over 72 hours normalized to body weight for Drd3-Cre::Htr4^loxP/loxP^ (cKO) mice and Htr4^loxP/loxP^ control littermates (WT) during the sucrose consumption test (SCT). Data were analyzed by repeated measures (RM) two-way ANOVA: time × genotype interaction, F(3,90) = 2.879, p = 0.0403; genotype factor, F(1,30) = 2.931, p = 0.0972 followed by post hoc Fisher’s LSD test, **p = 0.0077. n_WT_ = 16, n_cKO_ = 16. **b**, Quantification of water consumption during the SCT for each genotype. RM two-way ANOVA: time × genotype interaction, F(3,90) = 0.4320, p = 0.7306; genotype factor, F(1,30) = 0.09834, p = 0.7560. n_WT_ = 16, n_cKO_ = 16. **c**, Body weight of the mice of each genotype at the end of the SCT. Two-tailed unpaired t-test, p = 0.682. n_WT_ = 16, n_cKO_ = 16. **d**, Time spent in the center in the OF test over 60 min. Two-tailed unpaired t-test, p = 0.1443. n_WT_ = 20, n_cKO_ = 17. **e**, Quantification of locomotor activity (total distance traveled) for each 10-min bin in the OF test. RM two-way ANOVA: genotype factor, F(1,35) = 0.9715, p = 0.3311. n_WT_ = 20, n_cKO_ = 27. **f**, cKO mice showed a trend towards higher velocity (cm/s) in the EPM compared to WT. Two-tailed unpaired t-test, p = 0.0761. n_WT_ = 19, n_cKO_ = 16. **g**, Home cage food intake was measured for 30 min following the NSF. Two-tailed unpaired t-test, **p = 0.0044. n_WT_ = 20, n_cKO_ = 16. **h**, Quantification of time spent interacting with an unfamiliar mouse (Mouse) or a novel inanimate object (Object) in the three-chamber social interaction test for cKO or WT controls. Schematic of the testing apparatus is shown at left. RM two-way ANOVA: genotype factor, F(1,44) = 4.572, p = 0.0381 followed by post hoc Fisher’s LSD test, **p = 0.0038; ^n.s.^p = 0.9750. n_WT_ = 12, n_cKO_ = 12. **i**, cKO mice displayed larger acoustic startle responses (ASR) compared to WT to increasing levels of acoustic pulses. dB, decibel. RM two-way ANOVA: dB × genotype interaction, F(4,84) = 2.618, p = 0.0407; genotype factor, F(1,21) = 3.409, p = 0.0790 followed by post hoc Fisher’s LSD test, **p = 0.0012. n_WT_ = 11, n_cKO_ = 12. **j**, During the pre-pulse inhibition (PPI) test, cKO mice consistently exhibited higher startle responses to 120 dB acoustic stimuli preceded by any pre-pulse (PP) magnitude. RM two-way ANOVA: genotype factor, F(1,21) = 10.20, p = 0.0044 followed by post hoc Fisher’s LSD test, **p < 0.01, *p < 0.05. n_WT_ = 11, n_cKO_ = 12. **k**, cKO mice displayed similar levels of PPI as WT. RM two-way ANOVA: genotype factor, F(1,21) = 0.02151, p = 0.8848. n_WT_ = 11, n_cKO_ = 12. All data are presented as mean ± SEM.

**Fig. S4.**
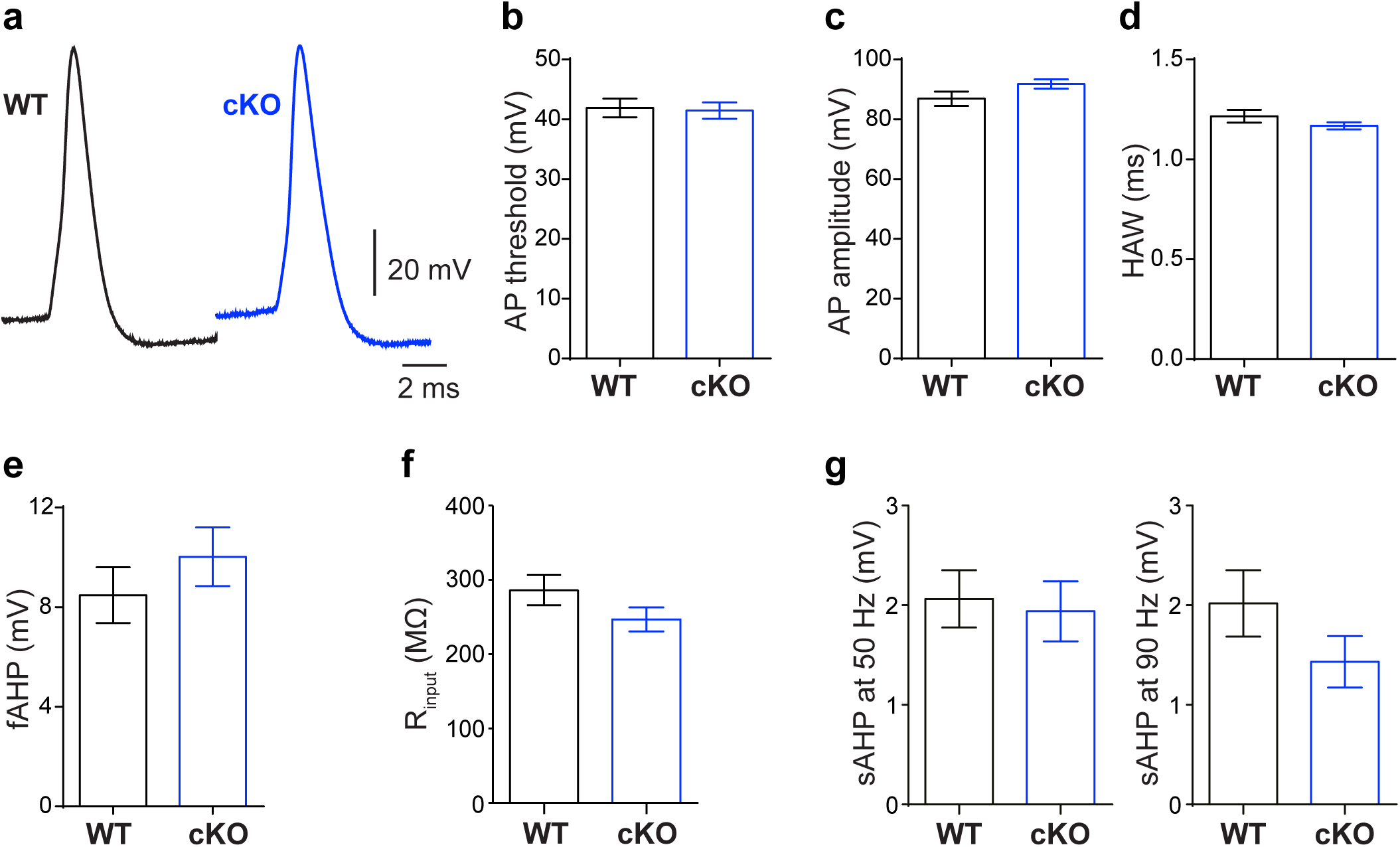
Effect of the loss of 5-HT_4_R on the baseline electrophysiological properties of DG GCs. Related to Fig. 3. **a**, Representative single action potentials (APs) from WT (black) and cKO (blue) DG GCs. **b-e**, No significant differences in AP properties were observed between genotypes: AP threshold (**b**); AP amplitude (**c**); HAW, half-amplitude width (**d**); fAHP, fast afterhyperpolarization potential (**e**). **f,** Input resistance (R_input_) was not different between genotypes. **g**, **h**, Slow afterhyperpolarization potential (sAHP) was similar between genotypes after both 50 Hz and 90 Hz trains of AP in 1 s. All data are presented as mean ± SEM. For both genotypes, 11 neurons from 3 mice were used for each experiment. Two-tailed unpaired t-test was used for all statistical analyses.

**Fig. S5.**
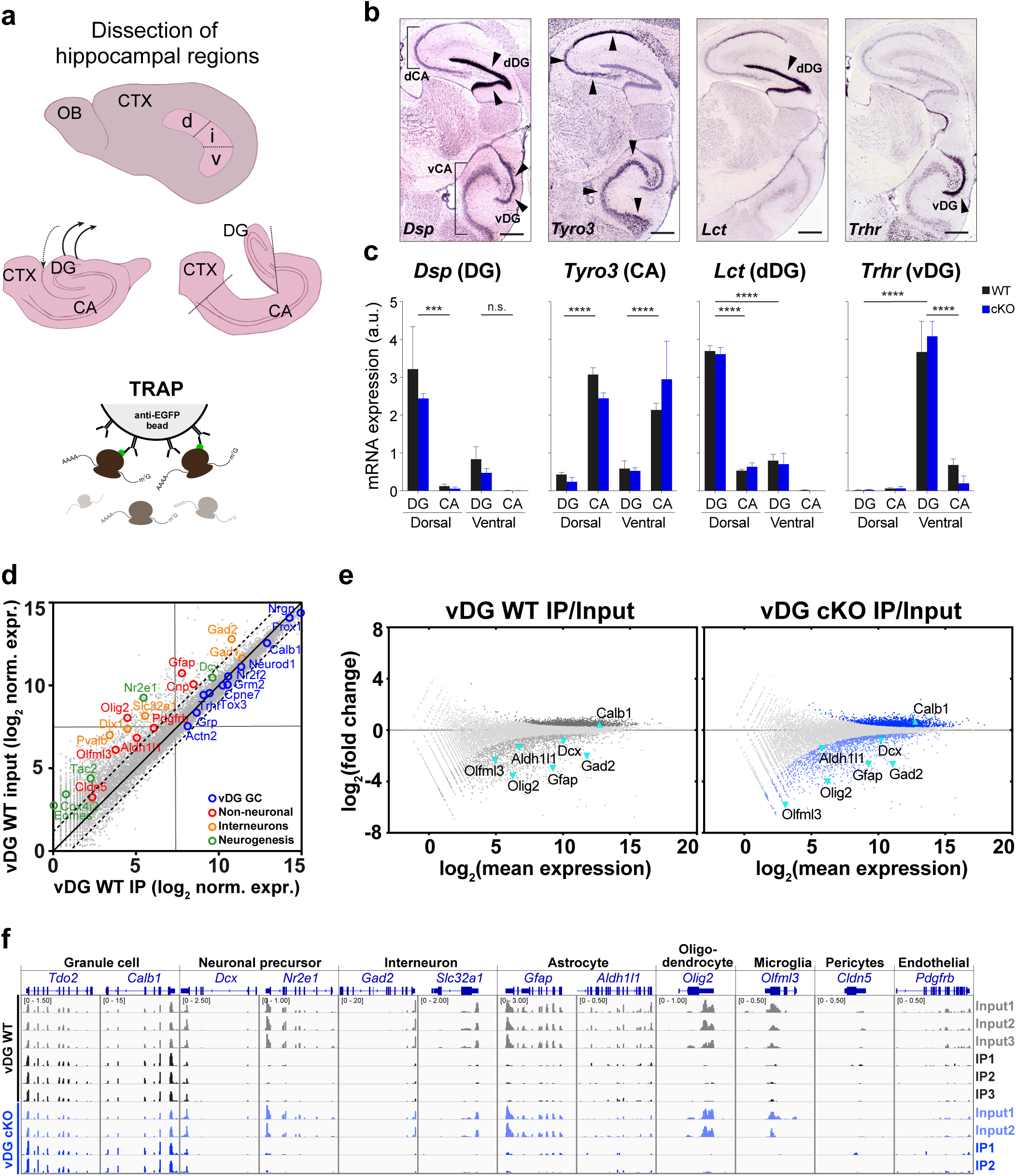
Assessment of regional and cell type specificity of vDG TRAP samples. Related to Fig. 4. **a**, Schematic representation of the dissection of hippocampal regions. **b**, In situ hybridization on sagittal sections through hippocampus showing the expression of region-specific genes. dDG: dorsal dentate gyrus, dCA: dorsal CA fields, vDG: ventral dentate gyrus, vCA: ventral CA fields. Arrowheads show region-specific expression for each gene. All in situ hybridization images in are © 2007 Allen Institute for Brain Science and available from: http://www.mouse.brain-map.org. **c,** qRT-PCR (mean ± SEM) showing relative expressions of region-specific genes from b in whole tissue mRNA samples. Two-way ANOVA followed by post hoc Fisher’s LSD test comparing region means. *Dsp:* genotype × region Interaction, F(3,12) = 0.2140, p=0.8848; genotype factor, F(1,12) = 0.6292, p = 0.4431; region factor, F(3,12) = 12.10, p = 0.0006; post hoc, ***p=0.0003, ^ns^p = 0.2501. *Tyro3*: genotype × region Interaction, F(3,12) = 1.916, p = 0.1809; genotype factor, F(1,12) = 0.005819, p=0.9405; region factor, F(3,12) = 34.25, p < 0.0001; post hoc, ****p < 0.0001. *Lct*: genotype × region interaction, F(3,12) = 0.2134, p = 0.8852; genotype factor, F(1,12) = 0.05478, p = 0.8189; region factor, F(3,12) = 279.0, p < 0.0001; post hoc, ****p < 0.0001. *Trhr*: genotype × region interaction, F(3,12) = 0.4318, p=0.7340; genotype factor, F(1,12) = 0.002429, p=0.9615; region factor, F(3,12) = 44.02, p < 0.0001; post hoc, ****p < 0.0001. **d**, Scatterplot comparing gene expression between input and IP in the vDG. Cell type marker genes are highlighted as vDG GC-specific (blue), non-neuronal (red), interneuron specific (yellow) and neurogenic (green). **e**, MA-plots showing differential expression analysis between IP and input samples from the vDG of WT (left) and cKO (right). Differentially expressed genes (FDR < 0.05) colored. Selected cell type marker genes are labeled. **f**, Genome browser view of RNA-seq reads mapped to selected cell type marker genes shows that IP samples, compared to input counterparts, were enriched in granule cell specific genes (*Tdo2*, *Calb1*), and depleted for genes specific for neuronal precursors (*Dcx*, *Nr2e1*), interneurons (*Gad2, Slc32a1*), astrocytes (Gfap, *Aldh1l1*), oligodendrocytes (Olig2), microglia (*Olfml3*), pericytes (*Cldn5*) and endothelial cells (*Pdgfrb*).

**Fig. S6.**
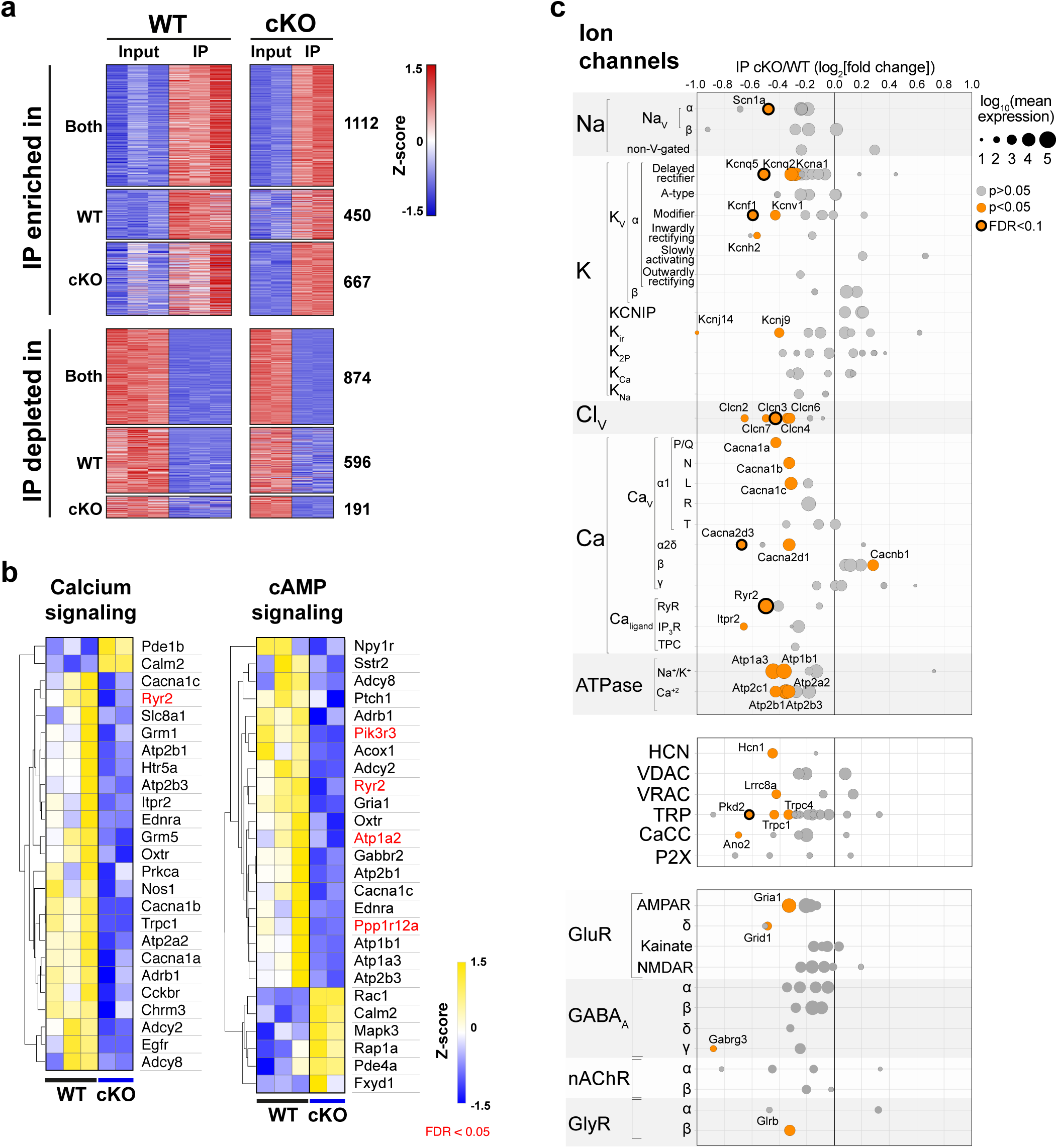
Genes related to 5-HT_4_R signaling and neuronal function were altered following 5-HT_4_R loss. Related to Fig. 4. **a**, Heatmap comparing normalized expression of IP enriched and depleted genes for each genotype in each sample. Detailed gene lists and values can be found in Table S2. **b**, Heatmap showing normalized expression values of calcium and cAMP signaling pathway-related genes in each TRAP IP mRNA from each genotype. Only genes with p < 0.05 are represented, red gene symbols indicate FDR < 0.05. **c**, Differential expression of ion channels in TRAP IP cKO/WT (log_2_[fold change]). Ion channels with normalized expression < 10 in the TRAP IP were excluded. Size of the bubbles are based on normalized mean expression of each gene across genotypes.

**Fig. S7.**
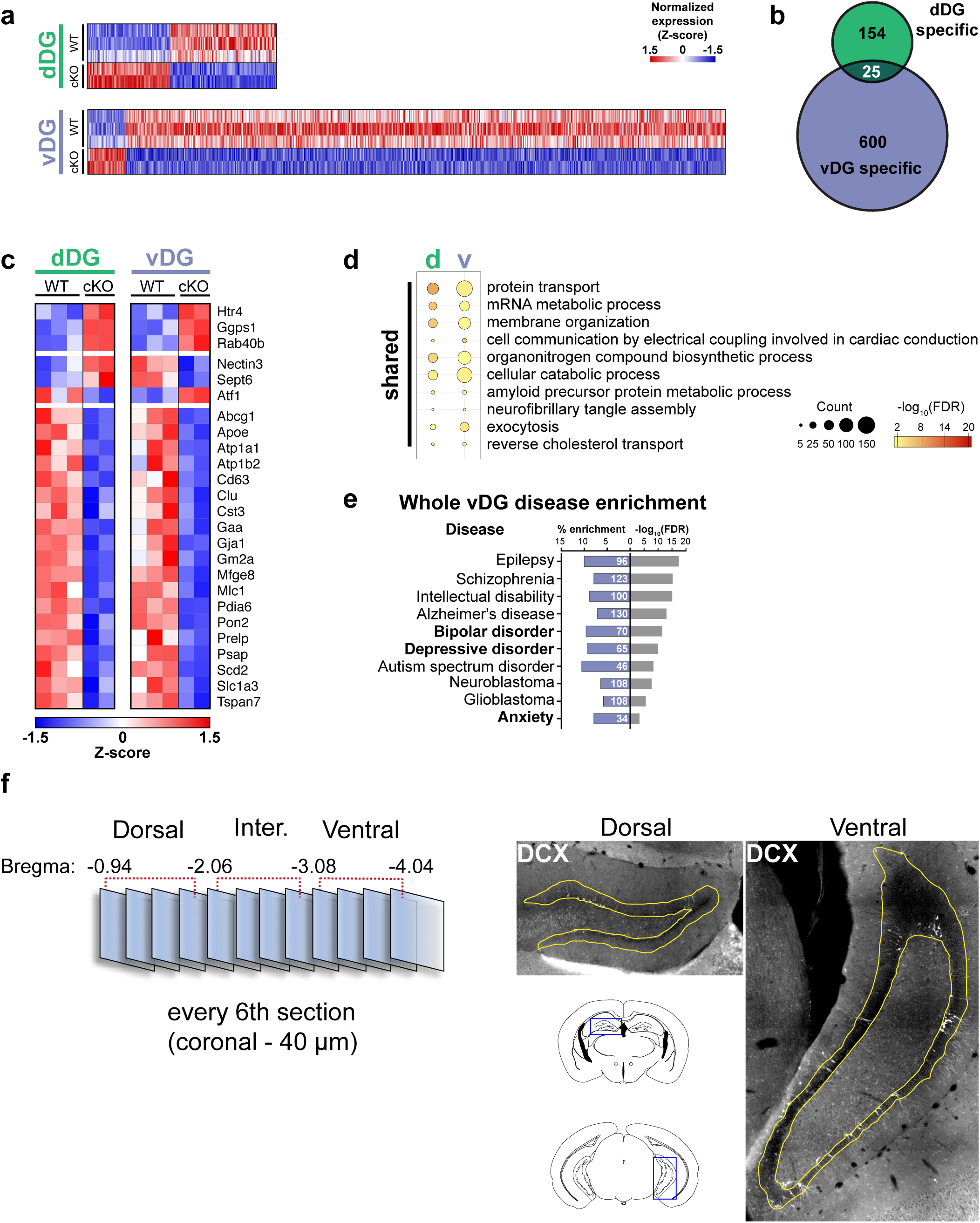
Comparison of the molecular properties of the dDG and vDG following 5-HT_4_R loss. Related to Fig. 5. **a**, Heatmaps of normalized expression values showing differentially expressed (DE) genes (FDR < 0.05) between WT and cKO in the whole tissue from dDG (top) and vDG (bottom). **b**, Venn diagram showing the number of overlapping DE genes between WT and cKO in the two regions. **c**, Heatmap showing 25 genes that were significantly altered in the cKO in both the dDG and vDG. Note that three genes (*Nectin3*, *Sept6,* and *Atf1*) were regulated in the opposite directions between two regions. **d**, Summary of shared gene ontology (GO) terms enriched in dDG and vDG (related to Fig. 5b). **e**, Summary of disease enrichment analysis of the DE genes in the vDG. For complete list, see Table S9. Note that bipolar disorder, depressive disorder and anxiety were among the significantly enriched diseases. No diseases showed significant association in DE genes of the dDG. **f**, Schematic depicting the experimental approach for quantification of neurogenesis. Left, the hippocampus was divided into three subdivisions (dorsal, intermediate, ventral) depending on the relative position of each coronal section to bregma. DCX+ cells were counted for every sixth 40 μm section. Right, example images of dorsal and ventral DG showing the area (highlighted in yellow) used for analysis. See **Materials and Methods** for details.

## Supplementary Material Legends

**Table S1. vDG IP cKO vs. WT DE genes.** Related to Fig. 4. List of differentially expressed (DE) genes between cKO and WT in vDG IP samples.

**Table S2. vDG IP vs. input enriched and depleted genes.** Related to Figs. 4 and S6. List of genes enriched and depleted in the IP compared to input for both WT vDG and cKO vDG samples.

**Table S3. vDG IP GSEA.** Related to Fig. 4. Gene set enrichment analysis results for vDG IP data sets.

**Table S4. vDG IP GO analysis.** Related to Fig. 4. Gene ontology analysis results for DE genes between genotypes in vDG IP samples.

**Table S5. dDG input cKO vs. WT DE genes.** Related to Figs. 5 and S7. List of DE genes between cKO and WT in dDG input samples.

**Table S6. vDG input cKO vs. WT DE genes.** Related to Figs. 5 and S7. List of DE genes between cKO and WT in vDG input samples.

**Table S7. dDG and vDG input GO analysis.** Related to Figs. 5 and S7. Gene ontology analysis results for DE genes between genotypes for both dDG input and vDG input.

**Table S8. vDG input only GO analysis.** Related to Figs. 5. Gene ontology analysis results for whole vDG (input)-specific DE genes.

**Table S9. vDG input disease enrichment.** Related to Figs. 5 and S7. Disease enrichment analysis results for DE genes between genotypes for vDG input.

**Document S1. *Htr4* Targeting Vector (TV).** Related to Figs. 1 and S1. Annotated TV sequence for the generation of the Htr4^Floxed^ line.

## Materials and Methods

### Animals

All procedures involving animals were approved by The Rockefeller University Institutional Animal Care and Use Committee and were in accordance with National Institutes of Health guidelines. Drd3-Cre (Drd3-Cre KI198) mice were generated by the GENSAT Project^1^ and were purchased from the Mutant Mouse Regional Resource Center (MMRRC) Repository (Stock #031741-UCD). Rosa26^fsTRAP^ mice were purchased from the Jackson Laboratories (Stock #022367). Htr4^Floxed^ and Htr4-bacTRAP mice were generated as described below. All animals were bred on a C57BL/6J background, group-housed, maintained on a 12-hr light-dark cycle at The Rockefeller University, and given ad libitum access to food and water. Animals used in the study were male, except for RNA sequencing experiments for which samples from male and female mice were pooled.

#### Generation of Htr4^Floxed^ mouse line

Targeting vector (TV) was designed in collaboration with inGenious Targeting Laboratory Inc., Ronkonkoma, NY. For annotated TV sequence (in full), see **Document S1**. A 7.92 Kb region used to construct the targeting vector (TV) was first subcloned from a positively identified C57BL/6 BAC clone (RP23:358G18) into a ∼2.4kb backbone vector (pSP72, Promega) containing an ampicillin selection cassette for retransformation of the construct prior to electroporation. A pGK-gb2 loxP/FRT neomycin resistance (Neo) cassette was inserted into the gene as depicted in **Fig. S1**. The region was designed such that the short homology arm (SA) extended 2347 base pairs (bp) 3’ to the Neo cassette. The long homology arm (LA) ended 5’ to the target region and was 4820 bp long. The loxP/FRT flanked Neo cassette was inserted 252 bp downstream of exon 5. The single loxP site, containing engineered MfeI and ApaI sites for southern blot analysis, was inserted 347 bp upstream of exon 5. The target region was 752 bp and included exon 5. The total size of the targeting construct (including vector backbone and Neo cassette) was 13.69 Kb. The targeting vector was confirmed by restriction analysis after each modification step, and finally, by sequencing with the following primers: P6 (5’-GAG TGC ACC ATA TGG ACA TAT TGT C-3’), T73 (5’-TAA TGC AGG TTA ACC TGG CTT ATC G-3’), N1 (5’-TGC GAG GCC AGA GGC CAC TTG TGT AGC-3’), and N2 (5’-TGC GAG GCC AGA GGC CAC TTG TGT AGC-3’). The P6 and T73 primers anneal to the backbone vector sequence and read into the 5’ and 3’ ends of the BAC sub-clone and the N1 and N2 primers anneal to the 5’ and 3’ ends of the loxP/FRT flanked Neo cassette and sequence the 5’ side of SA and 3’ side of the target region, respectively. The TV was linearized using Notl prior to electroporation into C57BL/6 mouse embryonic stem cells. Three lines of chimeric mice were generated at the Janelia Research Campus Gene Targeting and Transgenic Facility. One of these lines (1G12) showed germline transmission of the mutation. The 1G12 line was crossed to a germline Flpe recombinase line to remove the Neo cassette, leaving only two loxP sites flanking exon 5 of *Htr4*. We established this subsequent line, which are referred to as “Htr4^Floxed^ mice” in our laboratory at Rockefeller University.

#### Generation of Htr4-bacTRAP mouse line

To generate the Htr4-bacTRAP mice, the RP23-53D24 BAC, which contained the *Htr4* locus, was modified using the two-plasmid/one recombination protocol as described previously^2,3^. Briefly, a homology arm corresponding to the region immediately upstream of the ATG translation initiation site of the *Abcb1a* gene was cloned into the pS296 targeting vector^4^ containing EGFPL10a using the AscI and NotI restriction sites. Recombination was performed by electroporating the pS296-Htr4 vector into electrocompetent DH10β bacteria containing pSV1.RecA plasmid and the BAC. Successful recombination was determined by screening cointegrates by PCR and Southern blot analysis of HindIII digested BAC DNA, using the homology region as a probe. The modified BAC was prepared by double acetate purification with CsCl centrifugation followed by membrane dialysis and microinjected into the pronuclei of fertilized FVB/N mouse oocytes at a concentration of 0.5 ng/µl. Five transgenic founder mice were generated and crossed to C57BL/6J mice. F1 progeny were screened for proper transgene expression by EGFP immunohistochemistry. Founder line ES1299 showed accurate and robust expression of the transgene and was therefore selected for colony expansion.

#### Genotyping

At age 15-18 days, pups were marked with an ear punch for identification and tail biopsies were performed. Each tail sample was lysed for DNA extraction in 200 µl tail lysis buffer (100 mM Tris.HCl-pH8.8, 1 mM EDTA, 0.5% Tween20, 100 ug/mL proteinase K) at 56 °C for at least 24 hours. The tail DNA samples were analyzed via PCR using 1-2 µl of lysis solution, GoTaq DNA Polymerase (Promega) and corresponding primer pairs as listed below.

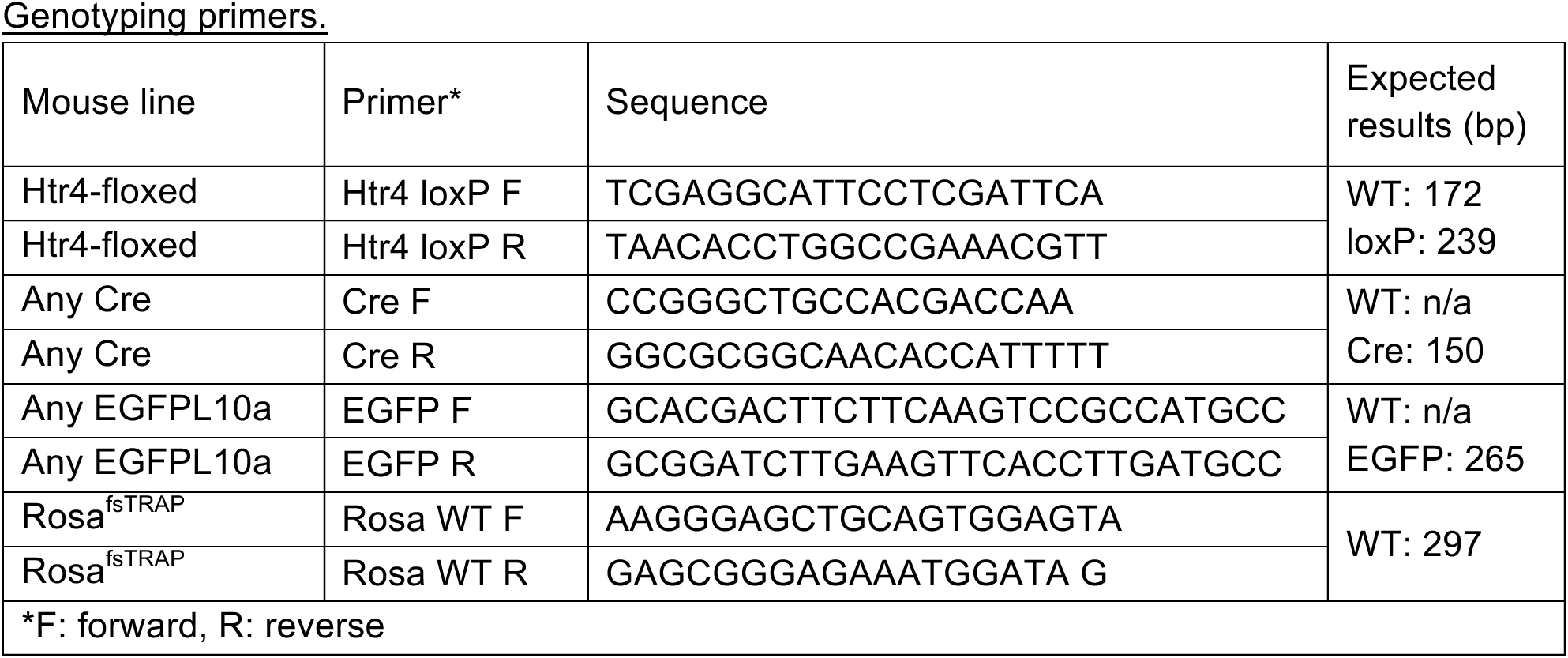

### cAMP induction assay

Intact *Htr4* (WT) and mutant *Htr4^delE5^* (delE5) cDNA coding sequences (CDSs) (inserts) were cloned into pCMV6-Entry-Myc-DDK tagged mammalian expression vector (OriGene, CAT#: PS100001) by homologous recombination-based In-fusion HD cloning kit (Clontech). The inserts were amplified by PCR using CloneAmp HiFi PCR Premix (Clontech) with primers “Htr4 infusion forward” (5’-AGA TCT GCC GCC GCG ATC GCC ATG GAC AAA CTT GAT GCT AAT GTG-3’) and “Htr4 infusion reverse” (5’-GCG GCC GCG TAC GCG TAG TAT CAC TGG GCT GAG CAG-3’) from the cDNA samples generated from the dentate gyrus of either Drd3-Cre::Htr4^loxP/loxP^ (cKO) mice for delE5, or Htr4^loxP/loxP^ mice for WT expression. The primers were designed using “Online In-Fusion Tools” (https://bit.ly/1eFKzsP) to include 5’ and 3’ homologous sequences to the ends of the linearized vector for In-Fusion HD cloning, and the *Htr4* CDS start codon, yet exclude the stop codon to allow for the expression of the MYC-DDK tag sequences. pCMV6-EGFP was used as the control. HEK293T cells (ATCC) cultured in 24-well plates (1.9 cm^2^ culture area, Corning) in Dulbecco’s Modified Eagle’s Medium (DMEM) (Gibco) supplemented with 10% fetal bovine serum (FBS) (Gibco) were transfected with the delE5, WT or EGFP plasmids using FuGENE 6 transfection reagent (Promega). Forty-eight hours after the transfection, all cells were incubated in stimulation buffer consisting of DMEM + 0.5mM IBMX (Sigma Aldrich) for 30 min. For cAMP induction, cells were further incubated in either stimulation buffer or 100µM Zacopride (Tocris Bioscience) in stimulation buffer for 45 min. The cells were then lysed with lysis buffer (0.1 M HCl, 0.1%Triton-X) for 20 min, and the lysates were centrifuged for 10 min at 21,500 x g. cAMP levels in the supernatants were measured by monoclonal anti-cAMP antibody based direct cAMP ELISA kit according to the manufacturer’s guidelines (Non-acetylated version, NewEast Biosciences). Four biological replicates and two technical replicates were performed for each experimental group. The cAMP level of each biological replicate was normalized to its mean protein concentration measured using Qubit protein assay kit (Invitrogen) and Qubit 3.0 Fluorometer (Invitrogen). Statistical analysis of normalized cAMP levels was performed in GraphPad Prism 7 using one-way ANOVA followed by Fisher’s LSD.

### Immunohistochemistry

For fixed brain serial slice preparations, mice were deeply anesthetized by ketamine/xylazine and transcardially perfused with 10 ml of phosphate buffered saline (PBS) followed by 30 ml of 4% paraformaldehyde (PFA) in PBS. Brains were post-fixed in 4% PFA overnight at 4°C and cryoprotected by 30% sucrose in PBS. Coronal sections (40 µm) were acquired using a freezing microtome (Leica SM2010 R Sliding Microtome). The sections were then transferred to PBS, or cryoprotection buffer (25% glycerol and 25% ethylene glycol in PBS, pH 7.4) for long term storage at −20°C. For both 3,3′-Diaminobenzidine (DAB) immunohistochemistry (IHC) and all immunofluorescence (IFC) procedures, free-floating serial sections were blocked for 60 min at room temperature in PBS containing 3-5% normal donkey serum (NDS, Jackson ImmunoResearch) or normal goat serum (NGS, Vector Laboratories) and 0.1% Triton X-100. Sections were then incubated in the same blocking buffer overnight at 4°C with primary antibodies listed below at corresponding dilutions. On the next day, the sections were washed three times with PBS each for 5 min, and then incubated for 2 h at room temperature with the appropriate Alexa dye-conjugated (for IFC) or horseradish peroxidase (HRPT)-conjugated (for DAB IHC) secondary antibodies (Invitrogen) diluted 1:500 in the blocking buffer. The sections were then washed three times with PBS each for 5 min. For IFC, the sections were counterstained with DAPI for nuclear staining at 1:10000 dilution followed by three times PBS wash each for 5 min. For DAB IHC, the staining was developed using SIGMAFAST™ DAB tablets (Sigma-Aldrich) following the manufacturer’s protocol. The sections were mounted on the Superfrost slides (VWR) and sealed with Prolong Gold mounting reagent (Invitrogen). The slides were imaged on a Zeiss LSM700 confocal microscope for Fig. 2a-d, S2a-d and 5e, or a Zeiss Axioskop 2 microscope for Fig1b, Fig S2b and Fig S7f. Brightness was optimized using ImageJ software post-acquisition.

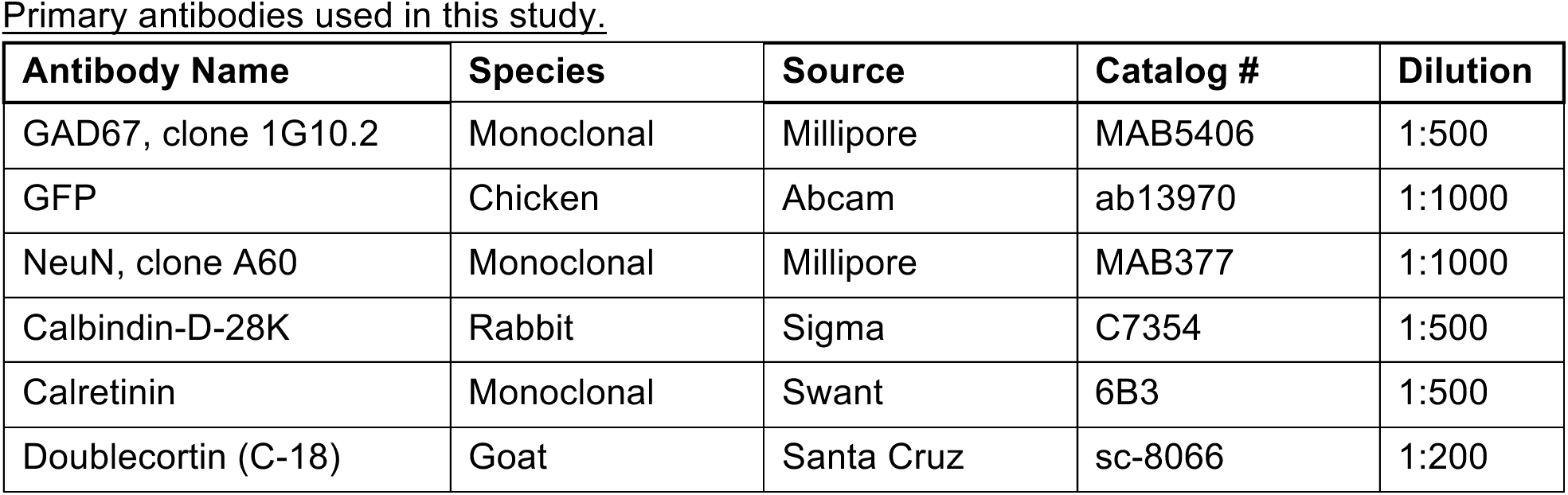

### Quantification of neurogenesis

An experimenter blinded to genotype counted number of DCX-positive (DCX+) cells in the DG granule cell layer (GCL) in every sixth 40 µm thick coronal sections through the entire hippocampus of WT or cKO mice (n = 5 per genotype). One hemisphere per animal was selected based on section quality and cells were counted at any depth through the entire section. Full size images of each section were then acquired and the area of GCL was calculated for each section by manually tracing the perimeter of the GCL as region of interest (ROI) in ImageJ. The hippocampus was divided into three subdivisions depending on the position of the coronal sections from bregma^5^: dorsal (−0.96 to −2.06), intermediate (−2.06 to −3.08) and ventral (−3.08 to −4.04). Each tissue section was registered to a subdivision (4 ± 1 sections per animal per subdivision). Neurogenesis was calculated as the number of DCX+ cells per area of GCL within a subdivision and values for each animal (cKO and WT) was normalized to the WT mean for each subdivision.

### Fluorescent in-situ hybridization (FISH)

Mice were transcardially perfused first in PBS and then 4% paraformaldehyde in PBS. After dissection, brains were post-fixed overnight at 4°C. They were then cryopreserved in 30% sucrose and sectioned using a cryostat (14 µm sections). Antigen retrieval was performed on fixed frozen tissue using an Oster Steamer, followed by FISH (RNAscope® Technology). The RNAscope® Multiplex Fluorescent Reagent Kit V2 (Advanced Cell Dagnostics) was used to target *Htr4* RNA using RNAscope® Probe Mm-Htr4 (Advanced Cell Diagnostics). Both antigen retrieval and FISH were performed according to manufacturer’s guidelines. Slides were coverslipped and imaged on an LSM700 confocal microscope.

### Total RNA isolation and quantitative RT-PCR (qRT-PCR)

Hippocampus tissue samples were dissected with fine tip forceps in ice-cold HBSS containing 2.5 mM HEPES-KOH (pH 7.4), 35 mM glucose, 4 mM NaHCO3 and total RNA was isolated using RNeasy Micro Kit (Qiagen) with on-column DNase digestion. RNA quantity was measured with a Nanodrop 1000 spectrophotometer (Thermo Scientific). cDNA samples were generated from 300-1000 ng of total RNA using qScript cDNA SuperMix (QuantaBio). All qRT-PCR experiments were performed on the LightCycler 480 System (Roche), using TaqMan assays (Applied Biosystems) and LightCycler 480 Probes Master mix (Roche). TaqMan assays used in this study are listed below. Default cycling conditions were followed (pre-incubation: one cycle, 95°C, 5 min; amplification: 45 cycles, 95°C for 10 s, 60°C for 30 s, 72°C for 1 s; ramp rate: 4.4°C/s). 10-20 ng of cDNA were used for each qRT-PCR reaction and three technical replicates were run for every sample. The mean C_T_ for each technical replicate was used for the quantification. Data were normalized to *Gapdh* as the endogenous control by the comparative C_T_ (2^-ΔΔCT^) method^6^.

| Gene Symbol | Assay ID | Dye |
| --- | --- | --- |
| Htr4 (3-4) | Mm00434129_m1 | FAM |
| Htr4 (4-5) | Mm01258807_m1 | FAM |
| Htr4 (5-6) | Mm01258808_m1 | FAM |
| Htr4 (6-7) | Mm01258809_m1 | FAM |
| Tyro3 | Mm00444547_m1 | FAM |
| Lct | Mm01285112_m1 | FAM |
| Dsp | Mm01351876_m1 | FAM |
| Trhr | Mm00443262_m1 | FAM |
| Gapdh | Mm99999915_g1 | FAM |

### Translating ribosome affinity purification (TRAP)

Affinity purification of EGFP-tagged polysomes was performed as previously described^7^, with minor modifications. Drd3-Cre::Rosa^fsTRAP^ (WT) and Drd3-Cre::Htr4^loxP/loxP^::Rosa^fsTRAP^ (cKO) mice were sacrificed, followed by rapid dissection of dorsal and ventral dentate gyrus with fine tip forceps in ice-cold HBSS containing 2.5 mM HEPES-KOH (pH 7.4), 35 mM glucose, 4 mM NaHCO3, and 100 μg/ml cycloheximide. Tissue from one male and one female mouse was pooled for each biological replicate. Pooled dentate gyrus samples were then homogenized in 1 ml extraction buffer containing 10 mM HEPES-KOH (pH 7.4), 150 mM KCl, 5 mM MgCl2, 0.5 mM DTT, 100 μg/ml cycloheximide, RNasin (Promega) and SUPERas-In (Life Technologies) RNase inhibitors, and Complete-EDTA-free protease inhibitors (Roche). Homogenates were cleared by centrifugation at 2000 x g at 4 °C. IGEPAL CA-630 (NP-40, Sigma) and DHPC (Avanti Polar Lipids) were both added to the supernatants (S2) to a final concentration of 1% for each, followed by centrifugation at 20,000 x g. Polysomes were immunoprecipitated from these supernatants (S20) using 100 μg monoclonal anti-EGFP antibodies (50 μg each of clones 19C8 and 19F7^4^) bound to biotinylated-Protein L (Pierce, Thermo Scientific) coated streptavidin-conjugated magnetic beads (Life Technologies), and washed in low salt buffer containing 10 mM HEPES-KOH (pH7.4), 150 mM KCl, 5 mM MgCl2, 1% IGEPAL CA-630, 0.5 mM DTT, 100 μg/ml cycloheximide, and RNasin RNase inhibitors (Promega). IPs were carried out overnight at 4°C and beads were washed with and washed in high salt buffer containing 10 mM HEPES-KOH (pH7.4), 350 mM KCl, 5 mM MgCl2, 1% IGEPAL CA-630, 0.5 mM DTT, 100 μg/ml cycloheximide, and RNasin RNase inhibitors (Promega). Bound RNA was purified using the RNeasy Micro Kit (Qiagen) with on-column DNase digestion. RNA was also purified from a fraction of the pre-IP S20 supernatant to serve as whole-tissue (input) samples. RNA quantity was determined using the Qubit RNA HS Assay kit (Invitrogen) and RNA quality was determined using Agilent 2100 Bioanalyzer with RNA 6000 Pico chips. Only samples with RNA integrity values ≥ 7.0 were used for RNA-seq.

### RNA-sequencing (RNA-seq)

For each sample, 15 ng of purified RNA was converted to cDNA and amplified using the Ovation RNA-Seq System V2 Kit (NuGEN) following manufacture’s guidelines. cDNA was fragmented to an average size of 250 bp using a Covaris C2 sonicator with the following parameters: intensity 5, duty cycle 10%, cycles per burst 200, treatment time 120 seconds. RNA-seq libraries were prepared from 10 μg amplified RNA using the TruSeq RNA Sample Preparation Kit v2 (Illumina) following manufacturer’s protocols and libraries were sequenced at The Rockefeller University Genomics Resource Center on the Illumina NextSeq 500 platform to obtain 75 bp paired-end reads. The RNA-seq datasets generated in this study are listed below.

Overview of RNA-seq datasets.
| Sample* | FastQ Name* |
| --- | --- |
| vDG_WT_IP_1 | Drd3-Cre_vDG_WT_IP_Rep1 |
| vDG_WT_IP_2 | Drd3-Cre_vDG_WT_IP_Rep2 |
| vDG_WT_IP_3 | Drd3-Cre_vDG_WT_IP_Rep3 |
| vDG_cKO_IP_1 | Drd3-Cre_vDG_Htr4_cKO_IP_Rep1 |
| vDG_cKO_IP_2 | Drd3-Cre_vDG_Htr4_cKO_IP_Rep2 |
| vDG_WT_Input_1 | Drd3-Cre_vDG_WT_input_Rep1 |
| vDG_WT_Input_2 | Drd3-Cre_vDG_WT_input_Rep2 |
| vDG_WT_Input_3 | Drd3-Cre_vDG_WT_input_Rep3 |
| vDG_cKO_Input_1 | Drd3-Cre_vDG_Htr4_cKO_input_Rep1 |
| vDG_cKO_Input_2 | Drd3-Cre_vDG_Htr4_cKO_input_Rep2 |
| dDG_WT_Input_1 | Drd3-Cre_dDG_WT_input_Rep1 |
| dDG_WT_Input_2 | Drd3-Cre_dDG_WT_input_Rep2 |
| dDG_WT_Input_3 | Drd3-Cre_dDG_WT_input_Rep3 |
| dDG_cKO_Input_1 | Drd3-Cre_dDG_Htr4_cKO_input_Rep1 |
| dDG_cKO_Input_2 | Drd3-Cre_dDG_Htr4_cKO_input_Rep2 |
| * brain region_genotype_RNA source_replicate |  |
| ‡ All data sets have been deposited in NCBI's Gene Expression Omnibus (GEO)<sup>8</sup>: accession number pending. |  |

### RNA-seq read mapping, analysis, and visualization

RNA-seq read quality was assessed by FastQC (0.11.4). Sequences were trimmed for trailing adaptors from the Illumina sequencing process by Trim Galor (0.4.1). The trimmed reads were then aligned to annotated exons using the mm10 mouse reference genome (UCSC) with STAR (2.4.2a)^9^ using default settings. SAMtools (v0.1.19-444) was used for indexing and removal of duplicates. The numbers of raw and mapped reads (bases), and the percentage of mapping for each sample were calculated via Picard (1.123) using the CollectRNASeqMetrics program and listed below. The quantification of aligned, sorted and indexed reads was done using the htseq-count module of the HTSeq framework (0.6.0)^10^, using the “union” mode with default settings to generate raw counts for each sample. Aligned bam files were converted to tdf format using igvtools (2.3.61) for Integrative Genomics Viewer (IGV) visualization. Differential expression analyses were performed using DESeq2^11^, R-package version 1.4.5. Differentially expressed (DE) genes for each analysis were determined based on adjusted p-values (p-adj), also termed as false discovery rate (FDR), and reported in Results or the figure legends. IP enriched genes were determined as IP over input fold change ≥ 1.25 with FDR < 0.05, and IP depleted genes as IP over input fold change < 0.5 with FDR < 0.05. For Fig. 4d-f, when determining DE genes between the cKO and WT TRAP mRNA, nine genes were detected as genes normally depleted in IP (IP depleted in WT) and excluded from further analysis (red genes in Table S2). For MA-plots, log_2_ of the fold change between genotypes (log_2_[fold change]) and log_2_ of the mean of the normalized expression values in all samples (log_2_[mean expression]) for each gene were used. Scatter plots were generated using normalized expression values (log_2_[norm. expr.]). Heatmap visualizations (one minus Pearson correlation) of normalized expression values were generated using web-based Morpheus software (Broad Institute, https://software.broadinstitute.org/morpheus) and z-scores were calculated using mean subtracted and SD normalized values by gene. Nominal p-values (p) were also given when normalized expressions of specific genes were reported. Full lists of DE genes can be found in Tables S1 (vDG IP cKO vs. WT), **S2** (IP vs. input for cKO and WT), **S5** (dDG input cKO vs. WT), and **S6** (vDG input cKO vs. WT). Gene set enrichment analysis (GSEA) was performed on the fold change-ranked gene list of IP samples comparing cKO and WT (p < 0.05, log[norm. expr.] > 1). We also excluded genes strongly depleted in the IP (WT IP/Input enrichment FDR < 0.05, log_2_[fold change] < −1) and *Htr4* as GSEA accounts for the direction of the regulation of gene expression and *Htr4* expression seemed upregulated although it was non-functional. GSEA desktop v3.0 software (Broad Institute) was used with the following parameters: Gene sets database = Molecular Signature Database (MSigDB), C2, Canonical Pathways (CP, c2.cp.v6.2.symbols.gmt); number of permutations = 1000; enrichment statistics = classic; set size, max = 100, min = 10; normalization = meandiv. Gene sets with FDR ≤ 25 are reported as advised^12^. Gene ontology (GO) and disease enrichment analyses were performed using ToppGene Suite (http://toppgene.cchmc.org)^13^. “GO: Biological Process” results were reported for GO analyses. FDR (Benjamini-Hochberg) < 0.05 was considered statistically significant. Network view in Fig. 4h was generated using the EnrichmentMap plugin (3.1.0) for Cytoscape (3.7.0) using enriched GO terms (FDR < 0.01, nodes) and the percentage of overlapping genes in among those terms (overlap > 50%, edges).

Quality control of RNA-seq alignments.
| Sample | Raw Reads | Mapped Reads | Ribosomal% | Coding % | UTR % | Intro-nic% | Interge-nic % | mRNA* % |
| --- | --- | --- | --- | --- | --- | --- | --- | --- |
| dDG_WT_Input_1 | 4976974951 | 4898777541 | 11.00 | 26.15 | 34.18 | 11.65 | 17.48 | 60.33 |
| dDG_WT_Input_2 | 4114194145 | 4048376030 | 9.60 | 28.97 | 36.85 | 9.12 | 15.87 | 65.82 |
| dDG_WT_Input_3 | 6348807839 | 6241867530 | 13.52 | 30.73 | 36.91 | 7.14 | 12.29 | 67.64 |
| dDG_KO_Input_1 | 5318465410 | 5237452256 | 9.40 | 29.55 | 37.17 | 8.97 | 15.33 | 66.72 |
| dDG_KO_Input_2 | 4494618359 | 4433172922 | 7.08 | 30.48 | 37.47 | 8.40 | 16.89 | 67.95 |
| vDG_WT_Input_1 | 5274266386 | 5200347558 | 7.32 | 29.76 | 37.30 | 9.69 | 16.25 | 67.07 |
| vDG_WT_Input_2 | 5908160994 | 5817160131 | 8.02 | 31.64 | 36.97 | 8.64 | 15.09 | 68.61 |
| vDG_WT_Input_3 | 5200024888 | 5110416182 | 12.45 | 30.12 | 36.81 | 7.70 | 13.47 | 66.93 |
| vDG_KO_Input_1 | 5390357255 | 5295752262 | 11.57 | 27.52 | 35.93 | 8.45 | 17.04 | 63.45 |
| vDG_KO_Input_2 | 5347253361 | 5264487834 | 10.03 | 29.78 | 38.46 | 8.79 | 13.38 | 68.24 |
| vDG_WT_IP_1 | 5055676715 | 4963214738 | 15.25 | 35.93 | 39.45 | 3.22 | 6.83 | 75.37 |
| vDG_WT_IP_2 | 5200875692 | 5096531166 | 18.77 | 34.24 | 37.98 | 2.97 | 6.88 | 72.22 |
| vDG_WT_IP_3 | 5176647321 | 5088146609 | 11.74 | 39.38 | 38.93 | 3.07 | 7.41 | 78.31 |
| vDG_KO_IP_1 | 4386834142 | 4294720920 | 18.98 | 33.93 | 37.81 | 3.40 | 6.72 | 71.74 |
| vDG_KO_IP_2 | 5041820503 | 4938661168 | 18.97 | 34.77 | 37.26 | 3.27 | 6.56 | 72.04 |
| * mRNA is the percentage of total bases mapping to Coding + UTR. |  |  |  |  |  |  |  |  |

### Electrophysiological recordings

Eight-week-old mice were euthanized with CO_2_. After decapitation and removal of the brains, transverse slices (400 μm thickness) were cut using a Vibratome 1000 Plus (Leica Microsystems) at 2 °C in a NMDG-containing cutting solution (in mM): 105 NMDG (N-Methyl-D-glucamine), 105 HCl, 2.5 KCl, 1.2 NaH_2_PO_4_, 26 NaHCO_3_, 25 Glucose, 10 MgSO_4_, 0.5 CaCl_2_, 5 L-Ascorbic Acid, 3 Sodium Pyruvate, 2 Thiourea (pH was around 7.4, with osmolarity of 295–305 mOsm). After cutting, slices were left to recover for 15 min in the same cutting solution at 35 °C and for 1 h at room temperature (RT) in recording solution (see below). Whole-cell patch-clamp recordings were performed with a Multiclamp 700B/Digidata1550A system (Molecular Devices). Dentate gyrus granule neurons were selected for recording based on their size, shape and position in the granular layer using an upright Olympus BX51WI microscope. The extracellular solution used for recordings contained (in mM): 125 NaCl, 25 NaHCO_3_, 2.5 KCl, 1.25 NaH_2_PO_4_, 2 CaCl_2_, 1 MgCl_2_ and 25 glucose (bubbled with 95% O_2_ and 5% CO_2_). The slice was placed in a recording chamber (RC-27L, Warner Instruments) and constantly perfused with oxygenated aCSF at 24 °C (TC-324B, Warner Instruments) at a rate of 1.5–2.0 ml/min. Whole-cell patch-clamp recordings were obtained from granule neurons using recording pipettes (King Precision Glass, Inc, Glass type 8250) pulled in a horizontal pipette puller (Narishige) to a resistance of 3–4 MΩ, filled with an internal solution containing (in mM): 126 K-gluconate, 4 NaCl, 1 MgSO_4_, 0.02 CaCl_2_, 0.1 BAPTA, 15 glucose, 5 HEPES, 3 ATP, 0.1 GTP (pH 7.3). For whole-cell recordings in the voltage clamp configuration, BIMU-8 (10 mM, Tocris Bioscience) was added to the bath for 1 minute and subsequently washed. Tetradotoxin (TTX, 1 mM, Tocris Bioscience) was added previously to the bath to avoid activation of neighboring neurons by BIMU-8. To measure the firing of the DG granule neurons, steps of 10 pA current were injected from a set starting membrane potential of −80mV. Data were acquired at a sampling frequency of 50 kHz and filtered at 1 kHz and analyzed offline using pClamp10 software (Molecular Devices).

### Behavior

Behavioral tests were performed on male mice from eight weeks up to four months of age. All mice were age-matched for each test, and control groups always consisted of WT littermates. For each test, mouse order was randomized, and the experimenter was blinded to genotype during the tests and data analyses. Mice were brought into the procedure room in their home cages and habituated to the room for one hour. Tests were performed within the light period of the light-dark cycle unless otherwise noted. The number of animals per group (n) and the statistical analyses are reported in the figure legends. For analysis of each behavioral tests, data points outside two standard-deviations of the mean for each group were determined as outliers and excluded. Statistical analyses were performed using GraphPad Prism 7 and 8 software, and p < 0.05 was considered significant.

#### Tail suspension test

Adhesive tape was used to suspend mice by their tail (0.5-1 cm from the tip of the tail) to the end of a piece of flexible plastic tubing hanging from the top center of one of the four compartments of a white acrylic box. Four mice were tested at the same time in each compartment and were separated by white acrylic walls. While suspending, mice were about 30 cm above the floor of the box. Test sessions lasted 6 min and were videotaped. Mice that held on to their bodies using their front limbs or started climbing up their tails were gently repositioned using a long stick without distracting other subjects. The floors and the walls of the box were cleaned with Clidox sterilizer and water in between trials. The amount of time spent immobile in the last 4 min of the test was quantified, which was extrapolated by scoring as mobile or immobile in every 5 s. In case of clasping to body parts or climbing up the tail, scoring was not applicable, and those mice were excluded from the analysis

#### Forced swim test

Mice were individually placed into glass cylinders (15 cm diameter, 35 cm height) filled with tap water (23-25 °C) to a height of 15.7 cm. Four mice were tested at the same time in separate cylinders placed as 2×2 adjacently. The cylinders were visually separated by white acrylic sheets. To start a trial, mice were gently placed into the cylinders keeping their head above the water surface. Test sessions lasted 6 min and were videotaped from above. The amount of time spent immobile, defined as the absence of all motions except floating and those required to keep a mouse’s head above the surface, was measured for the last 4 min of the trial, which was extrapolated by scoring as mobile or immobile in every 5 s.

#### Splash test

These experiments were performed in the dark cycle (20:00 – 23:59) to observe a higher amount of activity. The procedure room was illuminated with red light. Each mouse was sprayed with 200 µl 10% sucrose in tap water on their lower back using a 1 ml syringe with a needle tip attached. Immediately after, mice were placed in clean small mouse cages (22.2×30.80×16.24 cm) with bedding and filter top (no wire top). Four mice were tested at the same time in adjacent cages visually separated by white acrylic sheets. Test sessions lasted 5 min and were videotaped from the side. Grooming time was measured for a total 5 min, which was extrapolated by scoring as grooming or non-grooming in every 5 s.

#### Sucrose consumption test

Each mouse was placed in a clean cage with bedding and food, containing two water bottles, one filled with 1% sucrose in tap water and another filled with just tap water. Bottles were 50 ml Falcon conical tubes with sipper caps composed of 2.5 cm straight stainless steel ball-point sipper tubes (Ancare), inserted into rubber stoppers. Consumption of sucrose solution and water was monitored for 72 hours by measuring the decrease in the weight of the bottles every 12 h. At the each 12-hour mark, the bottles were switched to avoid side preference. At the end of the test, animals were weighed and amount of sucrose and water consumed (g) was normalized the weight per animal (g).

#### Open field test

Open field behavior was assayed in a square, acrylic arena (50×50×22.5 cm) with white floor and clear walls, equipped with two rows of infrared photocells placed 20 and 50 mm above the floor, spaced 31 mm apart. The procedure room was brightly and homogenously lit with fluorescent ceiling lamps. Each animal was placed in the open field arena for 60 min. Photocell beam interruptions were recorded on a computer using the Superflex software (Accuscan Instruments). The floors and the walls of the arenas were cleaned with Clidox sterilizer and water in between trials. The data was exported as time spent and distance traveled in total, peripheral and center areas in 10 min bins.

#### Elevated plus maze

The elevated plus maze consisted of two opposite open arms without sidewalls (35×5 cm) and two enclosed arms with black 14 cm high sidewalls (35×5×14 cm) connected by a common central platform (5 cm^2^). The entire plus-maze apparatus was elevated 30 cm off the ground. The experimental area was isolated from the experimenter with black curtains running down from the ceiling to the floor on four-sides and homogenously lit by ceiling mounted LED lamps on four sides, ensuring minimal shadows. The illumination was adjusted so the middle of each open arm was at 30-lux. Testing began by placing an animal on the central platform of the maze facing the same open arm. Trials lasted 5 min and recorded by a ceiling mounted camera connected to a computer. EthoVision XT 7.0 software (Noldus) was used to track the mouse and record the total time spent in each arm, total distance traveled, and velocity. The maze was wiped with 30% Ethanol in between trials. Mice that fell off the open arms were excluded from the analysis.

#### Novelty suppressed feeding

Mice were food deprived for 24-hour prior to testing. Body weight was monitored to determine efficiency of deprivation. Testing was performed in a brightly lit arena similar to the open field (50×50×22.5 cm) where a food pellet (1.8 cm length of 1.58×0.95 cm diameter oval pellets) was placed in the center on a circular filter paper (20 cm diameter, Fisherbrand). A white noise source was active throughout the habituation and testing. Four subjects were tested at the same time in adjacent arenas. The latency to bite the food within a 15 min trial was recoded. Following NSF, each animal was put in a clean vivarium cage for 30 min with food ad libidum after each trial. The weight of the food was measured before and after the 30 min test.

#### Acoustic startle response and pre-pulse inhibition

Startle and PPI testing were performed in a startle response systems (39×38×58 cm, SR-LAB system, San Diego Instruments) consisting of a non-restrictive clear Plexiglas cylinders (inner diameter 4 cm, length 13 cm) resting on a white Plexiglas platform and placed in a ventilated, sound-attenuated chamber. High frequency speakers, controlled by SR-LAB software, were mounted 33 cm above the cylinders and used to present all acoustic stimuli. Piezoelectric accelerometers mounted under the cylinders transduced movements of the animals which were digitized and stored by an interface and computer assembly. A dynamic calibration system was used to ensure comparable sensitivities across four separate chambers. The test mouse was placed into the startle chamber and allowed to acclimatize for 5 min. The time between each trial was 7-23 s. The trial consisted of four blocks. All sound intensities were presented in a pseudo-random order within a block. Beginning at startling stimulus onset, 65 consecutive 1-ms readings were recorded to obtain the peak amplitude of the animal’s startle response (Vmax). Five seconds after each stimulus, another 65 ms were measured which constitutes the no-stimulus trials. Background noise was 65 dB. In the first block (pre-test), mice were presented with 120 dB acoustic stimuli (65 ms) for five trials. Vmax were recorded per trial and averaged per subject. In the second block (ASR), acoustic stimuli were presented in five different intensities (120, 110, 100, 90 and 80 dB, 65 ms), each for four trials. Vmax was recorded for each trial and averaged for each stimulus intensity per subject. In the third block (PPI), a 120 dB acoustic stimuli (40 ms) was preceded by four different pre-pulse (PP) intensities (0, 3, 6, 12 dB, 20 ms) for ten trials. The time between the prepulse and the pulse was 100 ms. Vmax was recorded for each trial and averaged for each prepulse intensity per subject. PPI was calculated as the percent decrease in startle response in PP3, PP6 and PP12 compared to PP0 per subject. In the fourth block (post-test), 120 dB stimuli (65 ms) were presented for five trials. Vmax was recorded per trial, averaged per subject, and compared to the pre-test responses to assess the habituation of ASR during the testing.

#### Social interaction

Social behavior was measured using the three-chamber social interaction test. A rectangular test arena (58.5×43×22.7 cm) was made of white acrylic and two inner walls divided it equally into three chambers. Each inner wall had an open middle section with a removable sliding door, allowing free access between the middle and the two side chambers when the doors were removed. The social interaction test was comprised of two parts: a habituation session and a trial session. During the habituation session, two empty wire cups were placed in the left and right chambers. The test mouse was placed in the center chamber and allowed to explore all three chambers for 10 min. Next, the doors to the side chambers were closed, and the test mouse was placed in the middle chamber for the trial session. A stranger mouse of the same sex/appearance and a novel inanimate object (plastic building toy) of similar size were placed inside the left and right wire cups in a random order to balance for side preference. Doors between chambers were opened to allow the test mouse to explore the three chambers freely for 10 min. During all sessions, the arena was videotaped from above and the interaction time with each cup, defined as sniffing or touching cup, was manually measured from the recorded videos. Stranger mice were habituated to the wire cups for 10 min each day for three days prior to testing. The arena and the wire cups were thoroughly cleaned with Clidox sterilizer and water between subjects.

### Statistical analysis

All data analysis was performed using GraphPad Prism 7 and 8, Microsoft Excel or R^14^. Statistical parameters including the exact value of n, precision measures (mean ± SEM) and statistical significance are reported within the Results or the figure legends. Data were determined to be statistically significant when p < 0.05 by two-way ANOVA (ordinary or repeated measures [RM]) followed by post hoc Fisher’s LSD test, one-way ANOVA followed by post hoc Fisher’s LSD test, or two-tailed unpaired t-test. Statistical methods to analyze RNA-seq data are discussed in the “RNA-seq read mapping, analysis, and visualization”section.

## References

1. Ressler, K. J. & Mayberg, H. S. Targeting abnormal neural circuits in mood and anxiety disorders: from the laboratory to the clinic. Nat Neurosci 10, 1116–1124 (2007).

2. Price, J. L. & Drevets, W. C. Neural circuits underlying the pathophysiology of mood disorders. Trends Cogn. Sci. (Regul. Ed.) 16, 61–71 (2012).

3. Krishnan, V. & Nestler, E. J. The molecular neurobiology of depression. Nature 455, 894–902 (2008).

4. Ferguson, J. M. SSRI Antidepressant Medications: Adverse Effects and Tolerability. Prim Care Companion J Clin Psychiatry 3, 22–27 (2001).

5. Schmidt, E. F. et al. Identification of the cortical neurons that mediate antidepressant responses. Cell 149, 1152–1163 (2012).

6. Warner-Schmidt, J. L. et al. Cholinergic interneurons in the nucleus accumbens regulate depression-like behavior. Proc. Natl. Acad. Sci. U.S.A. 109, 11360–11365 (2012).

7. Bagot, R. C. et al. Circuit-wide Transcriptional Profiling Reveals Brain Region-Specific Gene Networks Regulating Depression Susceptibility. Neuron 90, 969–983 (2016).

8. Bagot, R. C. et al. Ketamine and Imipramine Reverse Transcriptional Signatures of Susceptibility and Induce Resilience-Specific Gene Expression Profiles. Biological Psychiatry 81, 285–295 (2017).

9. Hannon, J. & Hoyer, D. Molecular biology of 5-HT receptors. Behavioural Brain Research 195, 198–213 (2008).

10. Bockaert, J., Claeysen, S., Compan, V. & Dumuis, A. Neuropharmacology. Neuropharmacology 55, 922–931 (2008).

11. Andrade, R. in The Serotonin Receptors 481–494 (Humana Press, 2006). doi:10.1007/978-1-59745-080-5_16

12. Rosel, P. et al. Altered 5-HT2A and 5-HT4 Postsynaptic Receptors and Their Intracellular Signalling Systems IP_3_ and cAMP in Brains from Depressed Violent Suicide Victims. Neuropsychobiology 49, 189–195 (2004).

13. Marner, L. et al. Brain imaging of serotonin 4 receptors in humans with [11C]SB207145-PET. Neuroimage 50, 855–861 (2010).

14. Ohtsuki, T. et al. Association between serotonin 4 receptor gene polymorphisms and bipolar disorder in Japanese case-control samples and the NIMH Genetics Initiative Bipolar Pedigrees. Molecular Psychiatry 7, 954–961 (2002).

15. Lucas, G. et al. Serotonin4 (5-HT4) Receptor Agonists Are Putative Antidepressants with a Rapid Onset of Action. Neuron 55, 712–725 (2007).

16. Warner-Schmidt, J. L. et al. Role of p11 in Cellular and Behavioral Effects of 5-HT4 Receptor Stimulation. Journal of Neuroscience 29, 1937–1946 (2009).

17. Pascual-Brazo, J. et al. Modulation of neuroplasticity pathways and antidepressant-like behavioural responses following the short-term (3 and 7 days) administration of the 5-HT_₄_ receptor agonist RS67333. Int. J. Neuropsychopharm. 15, 631–643 (2012).

18. Mendez-David, I. et al. Rapid anxiolytic effects of a 5-HT_₄_ receptor agonist are mediated by a neurogenesis-independent mechanism. Neuropsychopharmacology 39, 1366–1378 (2014).

19. Faye, C. et al. Rapid anxiolytic effects of RS67333, a serotonin type 4 receptor agonist, and diazepam, a benzodiazepine, are mediated by projections from the prefrontal cortex to the dorsal raphe nucleus. BPS 1–60 (2019). doi:10.1016/j.biopsych.2019.08.009

20. Lucas, G. et al. Selective serotonin reuptake inhibitors potentiate the rapid antidepressant-like effects of serotonin4 receptor agonists in the rat. PLoS ONE 5, e9253 (2010).

21. Compan, V. Attenuated Response to Stress and Novelty and Hypersensitivity to Seizures in 5-HT4 Receptor Knock-Out Mice. Journal of Neuroscience 24, 412–419 (2004).

22. Eisch, A. J. & Petrik, D. Depression and Hippocampal Neurogenesis: A Road to Remission? Science 338, 72–75 (2012).

23. Bonaventure, P. et al. Mapping of serotonin 5-HT(4) receptor mRNA and ligand binding sites in the post-mortem human brain. Synapse 36, 35–46 (2000).

24. Hegde, S. S. & Eglen, R. M. Peripheral 5-HT4 receptors. FASEB J. 10, 1398–1407 (1996).

25. Tonini, M. & Pace, F. Drugs acting on serotonin receptors for the treatment of functional GI disorders. Dig Dis 24, 59–69 (2006).

26. Kaumann, A. J. & Levy, F. O. 5-hydroxytryptamine receptors in the human cardiovascular system. Pharmacology and Therapeutics 111, 674–706 (2006).

27. Anacker, C. & Hen, R. Adult hippocampal neurogenesis and cognitive flexibility — linking memory and mood. Nature Publishing Group 18, 335–346 (2017).

28. Tanaka, K. F., Samuels, B. A. & Hen, R. Serotonin receptor expression along the dorsal-ventral axis of mouse hippocampus. Philosophical Transactions of the Royal Society B: Biological Sciences 367, 2395–2401 (2012).

29. Gong, S. et al. Targeting Cre recombinase to specific neuron populations with bacterial artificial chromosome constructs. Journal of Neuroscience 27, 9817–9823 (2007).

30. Zhou, P. et al. Interrogating translational efficiency and lineage-specific transcriptomes using ribosome affinity purification. Proc. Natl. Acad. Sci. U.S.A. 110, 15395–15400 (2013).

31. Warner-Schmidt, J. Short-circuiting depression. Nature Publishing Group 19, 680–681 (2013).

32. O’Leary, O. F. & Cryan, J. F. in Mood and Anxiety Related Phenotypes in Mice: Characterization Using Behavioral Tests (ed. Gould, T. D.) 119–137 (Humana Press, 2009).

33. Hascoët, M. & Bourin, M. in Mood and Anxiety Related Phenotypes in Mice 42, 85–118 (Humana Press, Totowa, NJ, 2009).

34. Griebel, G. et al. Anxiolytic- and antidepressant-like effects of the non-peptide vasopressin V1b receptor antagonist, SSR149415, suggest an innovative approach for the treatment of stress-related disorders. Proc. Natl. Acad. Sci. U.S.A. 99, 6370–6375 (2002).

35. David, D. J. et al. Neurogenesis-Dependent and - Independent Effects of Fluoxetine in anAnimal Model of Anxiety/Depression. Neuron 62, 479–493 (2009).

36. Strekalova, T., Spanagel, R., Bartsch, D., Henn, F. A. & Gass, P. Stress-Induced Anhedonia in Mice is Associated with Deficits in Forced Swimming and Exploration. Neuropsychopharmacology 29, 2007–2017 (2004).

37. Gorman, J. M. Comorbid depression and anxiety spectrum disorders. Depress Anxiety 4, 160–168 (1996).

38. Kheirbek, M. A. et al. Differential Control of Learning and Anxiety along the Dorsoventral Axis of the Dentate Gyrus. Neuron 77, 955–968 (2013).

39. Gould, T. D., Dao, D. T. & Kovacsics, C. E. in Mood and Anxiety Related Phenotypes in Mice: Characterization Using Behavioral Tests (ed. Gould, T. D.) 1–20 (Humana Press, 2009).

40. Walf, A. A. & Frye, C. A. The use of the elevated plus maze as an assay of anxiety-related behavior in rodents. Nature Protocols 2, 322–328 (2007).

41. Samuels, B. A. & Hen, R. in Mood and Anxiety Related Phenotypes in Mice 63, 107–121 (Humana Press, 2011).

42. Crawley, K. R. B. A. J. N. Chapter 5: Anxiety-Related Behaviors in Mice. 1–25 (2008).

43. Miller, M. W. & Gronfier, C. Diurnal variation of the startle reflex in relation to HPA-axis activity in humans. Psychophysiology 43, 297–301 (2006).

44. Kaviani, H. et al. Affective modulation of the startle response in depression: influence of the severity of depression, anhedonia, and anxiety. J Affect Disord 83, 21–31 (2004).

45. Giakoumaki, S. G. et al. Low baseline startle and deficient affective startle modulation in remitted bipolar disorder patients and their unaffected siblings. Psychophysiology 85, 231–668 (2010).

46. Allen, N. B., Trinder, J. & Brennan, C. Affective startle modulation in clinical depression: preliminary findings. Biological Psychiatry 46, 542–550 (1999).

47. O’Brien-Simpson, L., Di Parsia, P., Simmons, J. G. & Allen, N. B. Recurrence of major depressive disorder is predicted by inhibited startle magnitude while recovered. J Affect Disord 112, 243–249 (2009).

48. Heiman, M. et al. A Translational Profiling Approach for the Molecular Characterization of CNS Cell Types. Cell 135, 738–748 (2008).

49. Heiman, M., Kulicke, R., Fenster, R. J., Greengard, P. & Heintz, N. Cell type-specific mRNA purification by translating ribosome affinity purification (TRAP). Nature Protocols 9, 1282–1291 (2014).

50. Vidal, R., Valdizán, E. M., Mostany, R., Pazos, Á. & Castro, E. Long-term treatment with fluoxetine induces desensitization of 5-HT4 receptor-dependent signalling and functionality in rat brain. J. Neurochem. 110, 1120– 1127 (2009).

51. Lucas, G. et al. Frontocortical 5-HT4 receptors exert positive feedback on serotonergic activity: Viral transfections, subacute and chronic treatments with 5-HT4 agonists. Biological Psychiatry 57, 918–925 (2005).

52. Castello, J. et al. CK2 regulates 5-HT4 receptor signaling and modulates depressive-like behavior. Molecular Psychiatry 23, 872–882 (2018).

53. Sargin, D. et al. Mapping the physiological and molecular markers of stress and SSRI antidepressant treatment in S100a10 corticostriatal neurons. Molecular Psychiatry 35, 395–18 (2019).

54. Andrade, R. & Chaput, Y. 5-Hydroxytryptamine4-like receptors mediate the slow excitatory response to serotonin in the rat hippocampus. J. Pharmacol. Exp. Ther. 257, 930–937 (1991).

55. Torres, G. E., Chaput, Y. & Andrade, R. Cyclic AMP and protein kinase A mediate 5-hydroxytryptamine type 4 receptor regulation of calcium-activated potassium current in adult hippocampal neurons. Mol. Pharmacol. 47, 191– 197 (1995).

56. Torres, G. E., Arfken, C. L. & Andrade, R. 5-Hydroxytryptamine4 receptors reduce afterhyperpolarization in hippocampus by inhibiting calcium-induced calcium release. Mol. Pharmacol. 50, 1316–1322 (1996).

57. Hoyer, D., Hannon, J. P. & Martin, G. R. Molecular, pharmacological and functional diversity of 5-HT receptors. Pharmacol. Biochem. Behav. 71, 533–554 (2002).

58. Barthet, G. et al. 5-hydroxytryptamine 4 receptor activation of the extracellular signal-regulated kinase pathway depends on Src activation but not on G protein or beta-arrestin signaling. Mol. Biol. Cell 18, 1979–1991 (2007).

59. Simms, B. A. & Zamponi, G. W. Neuronal voltage-gated calcium channels: structure, function, and dysfunction. Neuron 82, 24–45 (2014).

60. Thomas, G. M. & Huganir, R. L. MAPK cascade signalling and synaptic plasticity. Nat. Rev. Neurosci. 5, 173–183 (2004).

61. Medrihan, L. et al. Initiation of Behavioral Response to Antidepressants by Cholecystokinin Neurons of the Dentate Gyrus. Neuron 95, 564–576.e4 (2017).

62. Anacker, C. et al. Hippocampal neurogenesis confers stress resilience by inhibiting the ventral dentate gyrus. Nature Publishing Group 559, 1–25 (2018).

63. Samuels, B. A. et al. 5-HT1A receptors on mature dentate gyrus granule cells are critical for the antidepressant response. Nat Neurosci 18, 1606–1616 (2015).

64. Wang, X., Zhang, D. & Lu, X.-Y. Dentate gyrus-CA3 glutamate release/NMDA transmission mediates behavioral despair and antidepressant-like responses to leptin. Molecular Psychiatry 20, 509–519 (2015).

65. Kobayashi, K., Ikeda, Y., Haneda, E. & Suzuki, H. Chronic fluoxetine bidirectionally modulates potentiating effects of serotonin on the hippocampal mossy fiber synaptic transmission. Journal of Neuroscience 28, 6272–6280 (2008).

66. Costall, B. & Naylor, R. J. The influence of 5-HT2 and 5-HT4 receptor antagonists to modify drug induced disinhibitory effects in the mouse light/dark test. Br. J. Pharmacol. 122, 1105–1118 (1997).

67. Silvestre, J. S., Fernández, A. G. & Palacios, J. M. Effects of 5-HT4 receptor antagonists on rat behaviour in the elevated plus-maze test. European Journal of Pharmacology 309, 219–222 (1996).

68. Parfitt, G. M. et al. Bidirectional Control of Anxiety-Related Behaviors in Mice: Role of Inputs Arising from the Ventral Hippocampus to the Lateral Septum and Medial Prefrontal Cortex. Neuropsychopharmacology 42, 1715–1728 (2017).

69. Jimenez, J. C. et al. Anxiety Cells in a Hippocampal-Hypothalamic Circuit. Neuron 1–35 (2018). doi:10.1016/j.neuron.2018.01.016

70. Duric, V. & Duman, R. S. Depression and treatment response: dynamic interplay of signaling pathways and altered neural processes. Cell. Mol. Life Sci. 70, 39–53 (2013).

71. Pohodich, A. E. et al. Forniceal deep brain stimulation induces gene expression and splicing changes that promote neurogenesis and plasticity. Elife 7, R87 (2018).

72. Floriou-Servou, A., et al. Distinct Proteomic, Transcriptomic, and Epigenetic Stress Responses in Dorsal and Ventral Hippocampus. Biological Psychiatry 84, 531–541 (2018).

73. Gray, J. D. et al. Translational profiling of stress-induced neuroplasticity in the CA3 pyramidal neurons of BDNF Val66Met mice. Molecular Psychiatry 23, 904–913 (2018).

74. Duman, R. S. & Aghajanian, G. K. Synaptic Dysfunction in Depression: Potential Therapeutic Targets. Science 338, 68–72 (2012).

75. McEwen, B. S., Nasca, C. & Gray, J. D. Stress Effects on Neuronal Structure: Hippocampus, Amygdala, and Prefrontal Cortex. Neuropsychopharmacology 41, 3–23 (2016).

76. Santarelli, L. et al. Requirement of hippocampal neurogenesis for the behavioral effects of antidepressants. Science 301, 805–809 (2003).

77. Sahay, A. et al. Increasing adult hippocampal neurogenesis is sufficient to improve pattern separation. Nature 472, 466–470 (2011).

78. Strange, B. A., Witter, M. P., Lein, E. S. & Moser, E. I. Functional organization of the hippocampal longitudinal axis. Nature Publishing Group 15, 655–669 (2014).

79. Boldrini, M. et al. Antidepressants increase neural progenitor cells in the human hippocampus. Neuropsychopharmacology 34, 2376–2389 (2009).

80. Zhou, Q.-G., Lee, D., Ro, E. J. & Suh, H. Regional-specific effect of fluoxetine on rapidly dividing progenitors along the dorsoventral axis of the hippocampus. Sci. Rep. 6, 35572 (2016).

81. Wu, M. V. & Hen, R. Functional dissociation of adult-born neurons along the dorsoventral axis of the dentate gyrus. Hippocampus 24, 751–761 (2014).

82. Zheng, J. et al. Adult Hippocampal Neurogenesis along the Dorsoventral Axis Contributes Differentially to Environmental Enrichment Combined with Voluntary Exercise in Alleviating Chronic Inflammatory Pain in Mice. Journal of Neuroscience 37, 4145–4157 (2017).

83. Bielefeldt, A. Ø., Danborg, P. B. & Gøtzsche, P. C. Precursors to suicidality and violence on antidepressants: systematic review of trials in adult healthy volunteers. J R Soc Med 109, 381–392 (2016).

84. Birkett, M. A. et al. Acute anxiogenic-like effects of selective serotonin reuptake inhibitors are attenuated by the benzodiazepine diazepam in BALB/c mice. Pharmacol. Biochem. Behav. 98, 544–551 (2011).

85. Baek, I.-S., Park, J.-Y. & Han, P.-L. Chronic Antidepressant Treatment in Normal Mice Induces Anxiety and Impairs Stress-coping Ability. Exp Neurobiol 24, 156 (2015).

86. Kobayashi, K., Ikeda, Y. & Suzuki, H. Behavioral destabilization induced by the selective serotonin reuptake inhibitor fluoxetine. Mol Brain 4, 12 (2011).

87. Ansorge, M. S., Morelli, E. & Gingrich, J. A. Inhibition of serotonin but not norepinephrine transport during development produces delayed, persistent perturbations of emotional behaviors in mice. Journal of Neuroscience 28, 199–207 (2008).

## References for Materials and Methods

1. Gong, S. et al. Targeting Cre recombinase to specific neuron populations with bacterial artificial chromosome constructs. Journal of Neuroscience 27, 9817–9823 (2007).

2. Gong, S., Kus, L. & Heintz, N. Rapid bacterial artificial chromosome modification for large-scale mouse transgenesis. Nature Protocols 5, 1678–1696 (2010).

3. Gong, S., Yang, X. W., Li, C. & Heintz, N. Highly efficient modification of bacterial artificial chromosomes (BACs) using novel shuttle vectors containing the R6Kgamma origin of replication. Genome Res. 12, 1992–1998 (2002).

4. Heiman, M. et al. A Translational Profiling Approach for the Molecular Characterization of CNS Cell Types. Cell 135, 738–748 (2008).

5. Paxinos, G. & Franklin, K. B. J. The Mouse Brain in Stereotaxic Coordinates. (Academic Press, 2008).

6. Schmittgen, T. D. & Livak, K. J. Analyzing real-time PCR data by the comparative CT method. Nature Protocols 3, 1101–1108 (2008).

7. Heiman, M., Kulicke, R., Fenster, R. J., Greengard, P. & Heintz, N. Cell type-specific mRNA purification by translating ribosome affinity purification (TRAP). Nature Protocols 9, 1282–1291 (2014).

8. Barrett, T. et al. NCBI GEO: archive for functional genomics data sets--update. Nucleic Acids Res. 41, D991– 5 (2013).

9. Dobin, A. et al. STAR: ultrafast universal RNA-seq aligner. Bioinformatics 29, 15–21 (2013).

10. Anders, S., Pyl, P. T. & Huber, W. HTSeq--a Python framework to work with high-throughput sequencing data. Bioinformatics 31, 166–169 (2015).

11. Love, M. I., Huber, W. & Anders, S. Moderated estimation of fold change and dispersion for RNA-seq data with DESeq2. Genome Biol. 15, 550 (2014).

12. Subramanian, A. et al. Gene set enrichment analysis: a knowledge-based approach for interpreting genome-wide expression profiles. Proceedings of the National Academy of Sciences 102, 15545–15550 (2005).

13. Chen, J., Bardes, E. E., Aronow, B. J. & Jegga, A. G. ToppGene Suite for gene list enrichment analysis and candidate gene prioritization. Nucleic Acids Res. 37, W305–11 (2009).

14. R Core Team (2017). R: A language and environment for statistical computing. R Foundation for Statistical Computing, Vienna, Austria. http://www.R-project.org/.

