## Supplementary material for "Serotonin receptor 4 in mature excitatory hippocampal neurons modulates mood and anxiety": Document S1

### Construction of Conditional Targeting Vector for Htr4$^{Floxed}$

#### 1. Vector Design Outline

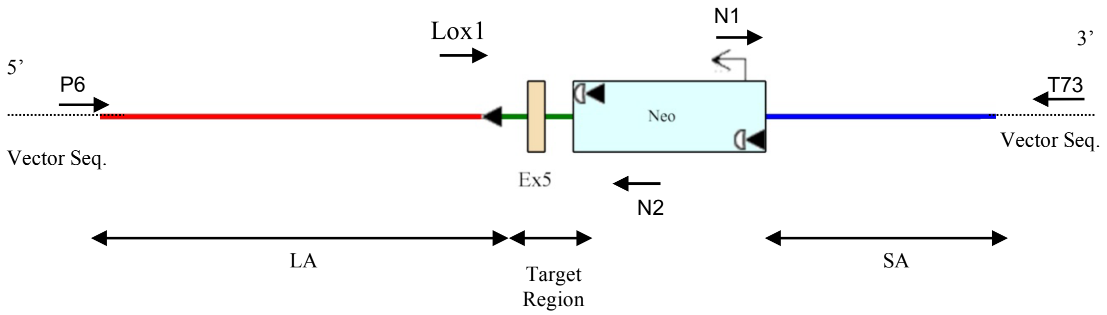


Ex5: *Htr4* exon 5
LA: Long arm
SA: Short arm
Neo: Neomycin resistance cassette
Black triangles: LoxP sites
White semicircles: FRT sites

#### 2. Sequence Data Analysis

#### PCR primers used for sequencing:

```
Primer P6:  5'- GAG TGC ACC ATA TGG ACA TAT TGT C   -3'
Primer T73: 5'- TAA TGC AGG TTA ACC TGG CTT ATC G   -3'
Primer N1:  5'- TGC GAG GCC AGA GGC CAC TTG TGT AGC -3'
Primer N2:  5'- TTC CTC GTG CTT TAC GGT ATC G       -3'
Primer lox1:5'- GGA CAG CAA TCT AAG GAG CAC AG      -3'
```

#### 3. Assembled Htr4$^{Floxed}$ targeting vector

1. Vector backbone: Blue text (e.g., ATCG)
2. LA (4820 bp): Bold and italic text (e.g., ***ATCG***)
3. Distal loxP (67 bp): Lightgray text, highlighted in yellow (e.g., ATCG)
4. MA (752 bp): Green text (e.g., ATCG)
5. Neo (1713 bp): Red text (e.g., ATCG)
6. SA (2347 bp): Underlined text (e.g., ATCG)
7. LoxP: Red text, highlighted in yellow (e.g., ATCG)
8. FRT: Red and underlined text, highlighted in gray (e.g., ATCG)
9. Exon: Highlighted in pink (e.g., ATCG)
10. Genotyping forward primer – Htr4 loxP F (**TCGAGGCATTCCTCGATTCA**)
11. Genotyping reverse primer – Htr4 loxP R (**TAACACCTGGCCGAAACGTT**, reverse transposed)

**3’end of BAC subclone joins here.**

CGATGATATCAGATCTGCCGGTCTCCCTATAGTGAGTCGTATTAATTTCGATAAGCCAGGTTAACCTGCAT
TAATGAATCGGCCAACGCGCGGGGAGAGGCGGTTTGCGTATTGGGCGCTCTTCCGCTTCCTCGCTCACTGA
CTCGCTGCGCTCGGTCGTTCGGCTGCGGCGAGCGGTATCAGCTCACTCAAAGGCGGTAATACGGTTATCCA
CAGAATCAGGGGATAACGCAGGAAAGAACATGTGAGCAAAAGGCCAGCAAAAGGCCAGGAACCGTAAAAAG
GCCGCGTTGCTGGCGTTTTTCCATAGGCTCCGCCCCCCTGACGAGCATCACAAAAATCGACGCTCAAGTCA
GAGGTGGCGAAACCCGACAGGACTATAAAGATACCAGGCGTTTCCCCCTGGAAGCTCCCTCGTGCGCTCTC
CTGTTCCGACCCTGCCGCTTACCGGATACCTGTCCGCCTTTCTCCCTTCGGGAAGCGTGGCGCTTTCTCAT
AGCTCACGCTGTAGGTATCTCAGTTCGGTGTAGGTCGTTCGCTCCAAGCTGGGCTGTGTGCACGAACCCCC
CGTTCAGCCCGACCGCTGCGCCTTATCCGGTAACTATCGTCTTGAGTCCAACCCGGTAAGACACGACTTAT
CGCCACTGGCAGCAGCCACTGGTAACAGGATTAGCAGAGCGAGGTATGTAGGCGGTGCTACAGAGTTCTTG
AAGTGGTGGCCTAACTACGGCTACACTAGAAGAACAGTATTTGGTATCTGCGCTCTGCTGAAGCCAGTTAC
CTTCGGAAAAAGAGTTGGTAGCTCTTGATCCGGCAAACAAACCACCGCTGGTAGCGGTGGTTTTTTTGTTT
GCAAGCAGCAGATTACGCGCAGAAAAAAAGGATCTCAAGAAGATCCTTTGATCTTTTCTACGGGGTCTGAC
GCTCAGTGGAACGAAAACTCACGTTAAGGGATTTTGGTCATGAGATTATCAAAAAGGATCTTCACCTAGAT
CCTTTTAAATTAAAAATGAAGTTTTAAATCAATCTAAAGTATATATGAGTAAACTTGGTCTGACAGTTACC
AATGCTTAATCAGTGAGGCACCTATCTCAGCGATCTGTCTATTTCGTTCATCCATAGTTGCCTGACTCCCC
GTCGTGTAGATAACTACGATACGGGAGGGCTTACCATCTGGCCCCAGTGCTGCAATGATACCGCGAGACCC
ACGCTCACCGGCTCCAGATTTATCAGCAATAAACCAGCCAGCCGGAAGGGCCGAGCGCAGAAGTGGTCCTG
CAACTTTATCCGCCTCCATCCAGTCTATTAATTGTTGCCGGGAAGCTAGAGTAAGTAGTTCGCCAGTTAAT
AGTTTGCGCAACGTTGTTGCCATTGCTACAGGCATCGTGGTGTCACGCTCGTCGTTTGGTATGGCTTCATT
CAGCTCCGGTTCCCAACGATCAAGGCGAGTTACATGATCCCCCATGTTGTGCAAAAAAGCGGTTAGCTCCT
TCGGTCCTCCGATCGTTGTCAGAAGTAAGTTGGCCGCAGTGTTATCACTCATGGTTATGGCAGCACTGCAT
AATTCTCTTACTGTCATGCCATCCGTAAGATGCTTTTCTGTGACTGGTGAGTACTCAACCAAGTCATTCTG
AGAATAGTGTATGCGGCGACCGAGTTGCTCTTGCCCGGCGTCAATACGGGATAATACCGCGCCACATAGCA
GAACTTTAAAAGTGCTCATCATTGGAAAACGTTCTTCGGGGCGAAAACTCTCAAGGATCTTACCGCTGTTG
AGATCCAGTTCGATGTAACCCACTCGTGCACCCAACTGATCTTCAGCATCTTTTACTTTCACCAGCGTTTC
TGGGTGAGCAAAAACAGGAAGGCAAAATGCCGCAAAAAAGGGAATAAGGGCGACACGGAAATGTTGAATAC
TCATACTCTTCCTTTTTCAATATTATTGAAGCATTTATCAGGGTTATTGTCTCATGAGCGGATACATATTT
GAATGTATTTAGAAAAATAAACAAATAGGGGTTCCGCGCACATTTCCCCGAAAAGTGCCACCTGACGTCTA
AGAAACCATTATTATCATGACATTAACCTATAAAAATAGGCGTATCACGAGGCCCTTTCGTCTCGCGCGTT
TCGGTGATGACGGTGAAAACCTCTGACACATGCAGCTCCCGGAGACGGTCACAGCTTGTCTGTAAGCGGAT
GCCGGGAGCAGACAAGCCCGTCAGGGCGCGTCAGCGGGTGTTGGCGGGTGTCGGGGCTGGCTTAACTATGC
GGCATCAGAGCAGATTGTACTGAGAGTGCACCATATGGACATATTGTCGTTAGAACGCGGCTACAATTAAT
ACATAACCTTATGTATCATACACATACGATTTAGGTGACACTATACCTGCAGGCGCGCCATTTAAATGCGG

CCGCTTAATTAACTCGACAAAAGCTGTTGGGATATGCCACTCCCCATGACAGGGAGGGTCTTCAGAGGGTA
AGACTAAAACTAAGCCCTGAATCATTAATTATGAACTAATGCATTTCAGCCTTTGATTTTCATTGTATATG
TTTTGTGTCTGTGTGGACCATTCATTACCTATAAGCATGGTAGCTCATCTTGGTCTGTTTTTTGGCCAGTT
TCTCCCTAATTAAGAATGCCAAAAGAACCAATCTGACCTCATCAACTTTGTGCTCTTGCTACAGACAAACA
AAGGAGCTGTCCCATACGTGTTTACATGAACATAGGCTGCTCTGAGATGTGCCTTGTTTTTTTATTACAAT
TTTGGTGTTAATTTTTCTTCTCTTCTGATAAATTTGACATAAGAAAAAAATGGGTAAGAGTGTGGCCTGGG
GTATTAGACTGCCTAGGTGCATTCTCACTTGAACAAATATGAGTTTCATCATCTTGAAAAAGCTACTTGAA
TTTTCTGGAGCTCACCTAGTTTGTAGAATTATGTTAATGACAGAAGCTTTCTCTGAGAGACTAAGGGAGTG
ACATAAATGAAGTGTGTGTGTGTGTGTGTGTGTGTGTGTGTGTGTGTGTGTGTGATGTATAAGCAGGTGCC
GCATACCTAATTCATCTCACTGATTTTAACCATTTGCCTTGGGAATTTTCCAAATACAAAGTGGCTTCTGC
AGGTGACTTCCTCGGCCTGACTCTTCAGCCTGATGAGCATCTGTTGTTGGAGAGTTAAGAGATTTCTCAGC
CTTGTGTGAAGCCTCCCCTTCCTTTGAAGAACTTACTTATCTGGTGGCTCACTTGAACCTTTCTTTTTGTT
CTCTCAGTTTACTTCTAGGCTCCAAGGTCCTCTCTCTATCTGTCTCTTATGAGGAGTACCTTCAGCATCTT
TCCTTTTCTCTCAAACCCAATAGGTTCAAAGTGGAAAATGTCCATCATACCTTAATAATCTTTTCTAGTTG
TTACATCAAAGGATTCCAAGTGTTGCCTTGTAGTTGCAGACATTGATACAAAATAGGACTTGCTTCTTGCC
ACAAAAGGTACTTCAATTAGCATGTGGCATATGCTGTAGATAATTGAAGTATCAGAGAAGAAAGCAGCAAT
TACAATGTAGTTGGAGAAACACCATGGAAACAGCAATGATACAGTGTGCTGAAGTGGGAAGCACATCCTTG
ATCTGGAGAGAAGGACACATTCAGAGAGCTGACCTAAAATATGGCTACTCCACAGCCTGGAGGAAGGGCTA
CTTCACTGCCCAGCTACACACACACACACACACACACACACACACACACACACACACAAATGTATAGGTAAAG
ATATAGGTATTGATATAGATGATATCTCTGTTTATCCTGTTGTAATTATGTGTGAGTATGGAAATTGACCC
CAAAATGTGTAAGCAATAGTATCTATTACATCTTATATGCAATGAACTCCATCATTCGCAGCCATTTACCT
AGAGGGAACAGCTTATTTGACGTTTTGTGTCTTAGACTCTCTAAGTACTAGGAGAGATGCTGATCCAGGCA
TTTTCATATGATGTATGTGCCAGGATCTTCCTATGTTTTCAACATATTATCTCATCTGTTCTCACAGCAAG
TGCATGGCGTAGCATCATTATCCCACTTTACAACTGAAGAAACTGTCTCGAAGAGGGTCAATAGCTAGCTC
TAAGTCTAGAGGGTATAAGAGTAAGGTAAAGGAAGTCAGTGGCCAACTGTGAGGGCACATGAGGTTGACTG
CTAGGTCTGCTAATTGTGAAAATCCAGGATTCCTGGTTGGTCTGGATGCTTCTCCTTTACCTATGGTTAGA
GAGCAGGGAATGAAAAGGTAAAATTTTCCTTAATGGCAGGAGTGGTGAAGACATTAGCTTGGGGCTTAAAA
TATACTGCTCTCTTCTAAGACTTGTCAAAGTAAATTGTGTCCCTGGAGACATTTAAAAGCAATTATGTCCC
TCATTTGCATGTGGCTGTGGTTGGAGAAGCTGCAGGTTGTCTACCAAAGGGGACTTTTATGGAGTTTTGTG
AGCAAACTTTGAAATGTTCAAAATAAGAACTTACTTTGTAAATTCCTTTGCTACATTCTAGCTAATCCTGG
CCATCTGGAACCAAATTTGCCATTACATTTATTGATTCATTTTGTAAAAGTATTTTTGCTTAACTAGAAAA
TGTCCTAGGAATGTCTAAAGAACAGAACTTATATAGTAGTATACTCCCAAGATAAACCACAGTTCTGTTTG
CTAAACAGCATCCTCTAGCCTGAGAATCAACAGCCCTCCTCTTTATTGGTTGTAATCCTCTCTGTTAGAAG
GGCATGCTGCAGGATTGACCTAGATGGGCATGTTCAAGTTCTGTGACCTCTTCTACATTAATAAACTCTTT
TAGAAGAATTACACCACTCCTTCTTTTCCTGCCTTGCATCTTTAGTTTTTGAGATGAGACTACCTTGAAAC
AGATGTCCTGGTCCTCTGGCTCTTACAATCTTTCTGGTGTCTCTTCCCCTGTGTCCCCTGAGCCTTAGATA
CAGGAATTGTGTTGTAGACCTATCCACTGGGGCTGGCACCCATGGCCAGCTCTTCTCTACAGTTTGAGCAG
TTGTTGTTTTGTGTTGGTCTCTATCTACTGAAAAAATGAAGCTCCTTCGATTCAGGGCAAAAACTACTCTT
ATCTGTAGGTGTAGGAACAGTACTTAGAATGCAGTTAGAAACCACACTGCTTTGAGAATGTGACAGTAAAT
TCTATGTCCCATGGCCTCGCCAGTTCTTATTTTTTCATCTTGAGACAGATACATGTGTGTATTTTGCCTTT
TAAAAAACTCAGGAGGGGCTGCTAGAAAGATATTCCAGTCACTAAAGTGACATGTTGTGCTCGTACAAGGA
CTTAAGTTCAAGCTTTAGAACTCATGTGAAAAGCTGGGCATTGTGGCACACACCTGTAATCTCAGGGTTGG
TGTTGGGTGTGGGGGTGGGAAGTGAGTAGACACAGGCAGGCCCCAGGGCTTGCCAGCCAGCCAGCTGAGAA
TATTTGATGAGTTCCTTTCCCATGAAAGACTCCTTTTCAAAAAAAAAAAAAAAAAACAGGTGAATGGCAATC
AAGATTGACCTCTGGCCGCCTCTGCACACATATAAACACAGGCTTGCCTACAAAACACACACACACACACA
CACACACACACACACACACTAGTTATGAGTGTTGTATATTCTAAGTACAGCCTTTAGTAAGGAAATCAGTG
GTAACGGTTACAATCAACTCTGTCAGTATATGTTAGGTGATCTCAGAACACTTGCCAATTTGACTTGCAAC
TTCTCACTAATGATGCTACCGTGCATATTTTAGAGGAAGGTCACAATACCATAAGGAAATTTTCTTAAAAA
CAAAACATACACAAAAAAACGTTTCTAGAATTTTTGCTTGAAATTGAAGATTTTATGATATGAGGAAAGGT
ACTCAAAACACATAGGTGGCTTTTTGAGAGTTCTGCCTTGAGAATACCAAAATCCTCAAAAGGAACCTTCA
GCATTTATGTCAATGTTGTGAGTTTTAGCTAAGAATCTCATATTTTGCCAGGCAGTGAGGCCAGTTTGGTC
TATAGACTGAATTTCAAGGCTACACAGAAAAACCCTGCCTTGAAATACAAAATACACAAACAAAAGCAAAA
ACCCTAATATTCTTTGTCCCATGCTACCAGTTAAAAATATTGTTATGTTTATTGTAGAATAAAGACAGAAT
TTAAGTCAAATATGAATCTGGGCATAATCACATATTTTCATTATATCTAAGCTAATCTAGCACTCCGGGTT
TCACCAACCTAGTTTAAAATTAAATAAAATAATCTCCTCCTCTTTTCTATTGCTAATGTTTTAAAGATGAG
GAAGAATTTTCCTATCTTCCTTAATAATTCATGTCTATGCAAGTTCAAATGTCCATAATTGTCTATACAGT
GGGCATCCCTGCCTCCCACACTAGCCCCTGACAACATCTTCAGTTACCCATTCCCCAGGATACTGACTGTA

***TCCAGAATATTTCCATTTGTTTCTCTGAATCAGCATCCTATACTAGTCTAGAGCAGTACCAATCAAGGCAT***
***AGAACATCTGGATAGACTGGAATAACATGGAGGATCACGTCTGGGAGTTGGAACTGAGGCCATTTGTGGAG***
***GAACAAGAAATTTAATACTTCAGGCAGAATATAAGAGGGTGAACAGTAAATTCCAAGAAAGGCTCCTATCA***
***CAGCAAATAAGGGAAGAGTACTTTCAACAAGCTGGCAAAATAACTAAGAGTAAGATGTTGGCAATTTTATA***
***TTCTTAGCTTTAGAGAAACAATTAAGCAATCACTGTCAGAAACAATGTATGTGAATTCTACTTGCCACATT***
***TCAGCTTGAGCATCTCATGTGCATGTACCCCTAGCCGGGAAATAAATAAGTGGTACTTCCTGGAGGTAGTG***
***TTGACTCAGGGGTGGACATGTACAGTCCAGGTGGCTGCCCGGCAAGCCGTGATACAGAAGACCCTCAGAAT***
***CTATGCTCTTCTCAGCTAAGTCTGTTGAGGGAGAAGAGAGGAGAAAGGGTAAGCCGGGACAGCAATCTAAG***
***GAGCACAGAAGTACACACAACATCTCAGCACCACCCTCAAGAAGAAGAGGACAAATCTGTGATAGTTCAAA***
***TTATCGTCACAGAAGCTAAAGGTGCCTGTGTGTAAGGGTCTGGAGAGCTCGAGGCATTCCTCGATTCACAG***
***AAAATGGGACTGTGTCCGAGGCAGAGTAGACCTTCATGCTACTTAAGACTTTGGGAATGAAGAGGGAAGTG***
***GAGCCAATT***GTATCACGCGTATAACTTCGTATAGCATACATTATACGAAGTTATGCCACTAGAGGATCCCC
GGGCCCGAGGCAATGTCCTTCACATGCACTATAAACGTATTTTATATAGTAACA**AACGTTTCGGCCAGGTG**
**TTA**ATGTTTTTATTGTATAATTGCCTTTATTATTCTCTCTGTTCTCTGCCCTCATCCCCTTCCTTTTCATT
TCCCTCCTCTGCCATCTCCTTCCCCACTTATTCCCAATTTTGTTTTCCCATTTTACTCTCCTCCTCTGATT
CTCCTTTTCTCAATATTTCCTCCTTCTTTCCATTTTTTTTTTACTCTTCTTGTCCTTCTCTCCCTTGTCCA
CTGTGGTCTTTCCCCTCTCCTTCTCATGTCCCCCTGCCTGTGGCCCCACCCTTAATTGTCTCTCCAGGTAT
TACGCCATCTGCTGCCAGCCTTTGGTTTATAGGAACAAGATGACCCCTCTACGCATCGCATTAATGTTGGG
AGGCTGCTGGGTCCTTCCCATGTTTATATCTTTTCTCCCCATAATGCAAGGCTGGAACAACATCGGCATAG
TTGATGTGGTAAGTATGCATACAAAGCCATGGGATTATTATCCACTCGGTTTTGTTAGGCATTTTAGATAA
GCAGACACTCTTAGCTAGGTAGCTATTTTAATAGATATTATAATAGTGGTCTGAGCAACATATTTAAATGT
CCTATTGATGCCTTGGTCAAATGTTAGTCAAGATCCTCTGAATTATCTTCCCAATGAGAACTTGTGTTCTT
AGAGTTCATTTATTTATTATTTGAAAATCTCATGCATAAGTGCAATCGTACGCCGGCTTAAGTGTACACGC
GTACTAGTCTAGCGAAGTTCCTATACTTTCTAGAGAATAGGAACTTCCCGCGGATAACTTCGTATAGCATA
CATTATACGAAGTTATGTAGATCTGATATCAGGGAGCTCTCAGACGTCGCTTGGTCGGTCTTTATTCGAAC
CCCAGAGTCCCGCTCAGAAGAACTCGTCAAGAAGGCGATAGAAGGCGATGCGCTGCGAATCGGGAGCGGCG
ATACCGTAAAGCACGAGGAAGCGGTCAGCCCATTCGCCGCCAAGCTCTTCAGCAATATCACGGGTAGCCAA
CGCTATGTCCTGATAGCGGTCCGCCACACCCAGCCGGCCACAGTCGATGAATCCAGAAAAGCGGCCATTTT
CCACCATGATATTCGGCAAGCAGGCATCGCCATGGGTCACGACGAGATCCTCGCCGTCGGGCATGCGCGCC
TTGAGCCTGGCGAACAGTTCGGCTGGCGCGAGCCCCTGATGCTCTTCGTCCAGATCATCCTGATCGACAAG
ACCGGCTTCCATCCGAGTACGTGCTCGCTCGATGCGATGTTTCGCTTGGTGGTCGAATGGGCAGGTAGCCG
GATCAAGCGTATGCAGCCGCCGCATTGCATCAGCCATGATGGATACTTTCTCGGCAGGAGCAAGGTGAGAT
GACAGGAGATCCTGCCCCGGCACTTCGCCCAATAGCAGCCAGTCCCTTCCCGCTTCAGTGACAACGTCGAG
CACAGCTGCGCAAGGAACGCCCGTCGTGGCCAGCCACGATAGCCGCGCTGCCTCGTCCTGCAGTTCATTCA
GGGCACCGGACAGGTCGGTCTTGACAAAAAGAACCGGGCGCCCCTGCGCTGACAGCCGGAACACGGCGGCA
TCAGAGCAGCCGATCGTCTGTTGTGCCCAGTCATAGCCGAATAGCCTCTCCACCCAAGCGGCCGGAGAACC
TGCGTGCAATCCATCTTGTTCAATGGCCGATCCCATGGTTTAGTTCCTCACCTTGTCGTATTATACTATGC
CGATATACTATGCCGATGATTAATTGTCAACACGTGCTGCTGCAGGTCGAAAGGCTCGGAGATGAGGAAGA
GGAGAACAGCGCGGCAGACGTGCGCTTTTGAAGCGTGCAGAATGCCGGGCCTCCGGAGGACCTTCGGGCGC
CCGCCCCGCCCCTGAGCCCGCCCCTGAGCCCGCCCCCGGACCCACCCCTTCCCAGCCTCTGAGCCCAGAAA
GCGAAGGAGCAAAGCTGCTATTGGCCGCTGCCCCAAAGGCCTACCCGCTTCCATTGCTCAGCGGTGCTGTC
CATCTGCACGAGACTAGTGAGACGTGCTACTTCCATTTGTCACGTCCTGCACGACGCGAGCTGCGGGGCGG
GGGGGAACTTCCTGACTAGGGGAGGAGTGGAAGGTGGCGCGAAGGGGCCACCAAAGAACGGAGCCGGTTGG
CGCCTACCGGTGGATGTGGAATGTGTGCGAGGCCAGAGGCCACTTGTGTAGCGCCAAGTGCCCAGCGGGGC
TGCTAAAGCGCATGCTCCAGACTGCCTTGGGAAAAGCGCCTCCCCTACCCGGTAGAATGAAGTTCCTATAC
TTTCTAGAGAATAGGAACTTCGTTCGAACATAACTTCGTATAGCATACATTATACGAAGTTATGGTACCTG
CAGAATTCATGCATAAGCTTGGATCCGTTCTTCGGACGCCTCGTCAACACCGTACGAATGTATTTAGATCA
CATGAAACCCTAAACAAGAAGTTACATTTCTGTGTCTGTCTTAGATTCTTTTCCTGTGCAATAATAAAATG
CCCTGACAGAAACAACTTAAGGGAGAAATGGCTTATTTTAGCTCACAGTTTAAGATATCCATCATGGCAGA
GATGTCAAGAGAGCAGGGACCTGCAGCAGTGAGTCCCTTAATGTCCATAGTCAGAAGACAATTAAACTGCT
CAGTTTGGGGTTTGTTTGTTTGTTTGTTTGTTTTGGGTTTGTTTGTTTTGGCTTCTACATTCTAGAACCCC
GTGTTAGGGTAAGGTCCTACCCACAACTAAGATGGATCTTCCCACACCAATTACCATACACAAGATAATCT
TCCCCGCTATACCCAGTGGCACTTCTTGTAGGTGATTCAGACTCTGTCAAGTTGACAATTCCCTCTAACCA
ACACAATGTCCAACTTTGCATTGCTCAGTTTCAGCAAGGATGCTGATGAGTCGGTTTATCTTGAATCCCCC
AGGACCAGGCACTTCATCCTTCACCTTTCCCATGAGATGTCTGATTGCCCTGGCTTGTCTTCAGCAAGAAT
CCTGCTAAGTTGACTTTCCCACTTTCTCTTCCCATTTACTCTGAATAGTTTTCTGTTCACTGACCCTCATC
CATCAATTCCTTGGCTATACATTCCCCACCACCACCACCACCTTTTTTTCCCTTTGGTGTTTTAGAGTTGA

ATCTAATCTCTTTGTGAAAAAAATCACAGTATTTTCTCTTATGAAAATAATGTGAAATATGTGGAATAAAG
GCTCCTTAGCAGGTGTTTCAAATATTTTTTTAACATTACAGTGAATCAGACATTACAATAAGGAATGACAT
TACAATAAGGAATTATACTTTTGGAAAGAGTATAATTATGGCAGTTTAAAGATGTGCAGGTTAATTTGCAC
TTCCCATGACACATAGTTTTCCTATAAAAAGAGTTTTAATTTCATTTATTTTATATATTTATTAAATATTT
ATTGATATTTATAAATCAGTTTCCAGAACTTGAGGCATATCATGTCCCAAGAAGTCTTCGTTATATTTCAT
ATTGAGTCTATAAGGATTACAGTGCCTATGGAAAGGCATACCAAGTAAATGGACAATGCCAAGGAATATGA
AAATCCTGGGCCAGAAGTCAGGAACTAAGCTGTGAGCCCAGCTCTGCTAATGCCCAACTTCCTGAAGAACT
GTATTAAGAAGGGAAGTTGGTGCTGAGTGAAAAATTGTCTAGTGTGAGTCTAAGGGGACCCATGCCACATA
GGCCATACATTGTCCAAACTCTGGATATTGTTTTCCAGAGACATAGAACTAATAGGATGTTTGTGTGCATA
TGTTCATCTATCTGTCTATCCATCTATATGTCTATCCACCTATCTGTCCATGAATCTATCTGTCATCTATC
TGTCTGTCTGTCTAACCATCTATCTACTATCTATTATCTATCTAACTATCTATCTGTCTGTCTGTCTGTCT
AACCAACCATCTATCTACTATCTATTATCTATATATCTACATTTCTATTTATCTTCTAATCTATCTAATCT
ATCTATCTATCTATCTATCTATCTATCTATCTATCTATCTATCATCTTTTCCTTGTCATGTATCTCTTTAT
CTATTTACCTATACACTTAGAAAAAGAAAAAGAGAAACAGAGGCAGACATATAGACATATTGTAACTGGTC
CTCCTGGTTGTGGAAGTTGATAATTCTCAGCAAACTAGAGACTGAGGAAAGGCAATATTTCAACTCAAGTC
CAAATATAGGAAAAAACTGATGGCCTTCATTAAGTCAATAGAGCAGTTCCCTCTTATTCGGTTATTTTGTT
CTATTTGGGCCTTTAATAGAGTTAGATGCAGTGCTAAATCGGAAGAGAAAATGGATTTATACAATCTGCCA
GTACAGAGCTTAATCTCAATCCCAAATGTTGTCGTACACAAAATTGGGTTTTACCAATTATCTGGATACCT
ATGACTTGGTTACACTGGCACATAACATTAACCATCTTGCTGTCAAAATCATCTACTCTGGCACACCAAAT
TATAATGATGATTAAAACTGATAGTCCAGATTCATTGTCAGGGCTATACATTTCTAAAGGAATATGGCATG
CAGTCACTTATAGCCAGTCCTAAGATCTTGCTAGATGTGGAGAAGGGGGCAGAAAGAAGAGGTTGAGAATG
AAGAATGAGTTCTGACCCAAGTGGATCAGAAAAAAAGAGAATTTGCCTTCTTTTCTGCACACCTAAGGACT
TTAATTTTATTTTCAAATGAGTCCCATGGCCAGGACATTTCTGAGTA

**5’ end of BAC subclone joins here.**
